# Leveraging gene correlations in single cell transcriptomic data

**DOI:** 10.1101/2023.03.14.532643

**Authors:** Kai Silkwood, Emmanuel Dollinger, Josh Gervin, Scott Atwood, Qing Nie, Arthur D. Lander

## Abstract

**BACKGROUND:** Many approaches have been developed to overcome technical noise in single cell RNA-sequencing (scRNAseq). As researchers dig deeper into data—looking for rare cell types, subtleties of cell states, and details of gene regulatory networks—there is a growing need for algorithms with controllable accuracy and fewer *ad hoc* parameters and thresholds. Impeding this goal is the fact that an appropriate null distribution for scRNAseq cannot simply be extracted from data when ground truth about biological variation is unknown (i.e., usually).

**RESULTS:** We approach this problem analytically, assuming that scRNAseq data reflect only cell heterogeneity (what we seek to characterize), transcriptional noise (temporal fluctuations randomly distributed across cells), and sampling error (i.e., Poisson noise). We analyze scRNAseq data without normalization—a step that skews distributions, particularly for sparse data—and calculate *p*-values associated with key statistics. We develop an improved method for selecting features for cell clustering and identifying gene-gene correlations, both positive and negative. Using simulated data, we show that this method, which we call BigSur (<u>B</u>asic Informatics and <u>G</u>ene <u>S</u>tatistics from <u>U</u>nnormalized <u>R</u>eads), captures even weak yet significant correlation structures in scRNAseq data. Applying BigSur to data from a clonal human melanoma cell line, we identify thousands of correlations that, when clustered without supervision into gene communities, align with known cellular components and biological processes, and highlight potentially novel cell biological relationships.

**CONCLUSIONS:** New insights into functionally relevant gene regulatory networks can be obtained using a statistically grounded approach to the identification of gene-gene correlations.

## Background

Single cell RNA-sequencing (scRNAseq), along with the related method of single nucleus RNA-sequencing, now offers researchers unparalleled opportunities to interrogate cells as individuals. Methods have been developed to classify cell types; identify gene expression markers; infer lineages; learn gene regulatory relationships, and examine the effects of experimental manipulations on both levels of gene expression and cell type abundances [1–7]. Because scRNAseq data are noisy, reliable inference requires leveraging information across many cells, trading off sensitivity for statistical power. How to handle that tradeoff should depend, ideally, on one’s goal. Simply clustering large numbers of transcriptionally very different cells (“cell types”) into small numbers of groups of similar size allows for a great deal of latitude in aggregating information across cells; it is thus not surprising that many different clustering approaches perform well. Other tasks, such as ordering cells along a continuum of gene expression change, or picking out rare cell populations within much larger groups, are less forgiving, and a plethora of different approaches currently compete for investigators’ attention [8–23]. Assessing the performance of such methods is frequently hindered by a lack of knowledge of ground truth.

A particularly challenging application of scRNAseq is the identification of patterns of gene co-expression. The identification of large-scale blocks of co-expressed genes—co-expression “modules”—can provide an alternative method for classifying cells when traditional clustering fails [24]. In contrast, smaller-sized blocks of gene co-expression have the potential to reflect true gene-regulatory networks that relate to specific functions [25]. This is because random transcriptional noise in gene circuits should induce weak but real correlations among regulatory genes and their targets. Indeed, it has long been proposed that gene regulatory links could be discovered solely from the weak gene expression correlations that one might encounter when studying otherwise homogenous populations of cells [26–32].

Unfortunately, identifying small yet significant gene expression correlations in single cell data requires a degree of statistical power that scRNAseq applications rarely strive for (and, to be fair, rarely need to). Yet, as greater numbers of scRNAseq datasets accumulate, with a growing trend toward increasing numbers of cells per dataset, we wondered whether substantial amounts of novel information about gene co-regulation might be accessible simply through a more in-depth examination of pairwise gene expression correlations.

One of the main challenges in pursuing such a program is the absence of an accepted statistical model for pairwise correlations in scRNAseq data. Only with a model can one define a null hypothesis by which to judge whether observations are significant. Unfortunately, with scRNAseq, there is not good consensus regarding the model to use for the data distributions of individual genes, much less their correlations. The common approach of fitting individual gene data to *ad hoc* analytical distributions (e.g. “zero-inflated negative binomial” [33]; reviewed by [34]), has met with frequent criticism that is difficult to dismiss [35–38]. One may seek to circumvent such concerns by attempting to learn empirical distributions on a case-by-case basis from data, but this typically requires making assumptions about the amount and distribution of actual biological variation in the data, which are frequently unknown. Furthermore, pitfalls in implementing empirical methods can be hard to avoid, particularly with high-order statistical information, such as correlations. For example, the seemingly reasonable intuition that one might be able to construct the distribution of the correlation coefficient under the null hypothesis simply by randomly permuting elements is actually incorrect [39].

One of the major obstacles to defining an appropriate data distribution for scRNAseq data is the fact that underlying sources of technical variation are not fully understood, nor is the range of biological variation in biologically “equivalent” cells fully known. Here we begin by re-considering these factors, and leveraging the work of others, in pursuit of an analytical model of null correlation distributions that makes the fewest ad hoc assumptions and minimizes adjustable parameters. We show that the approach that emerges has the power to identify subtle yet real correlations, both positive and negative, in scRNAseq data, even among genes in modest numbers of cells that are relatively sparsely sequenced. Ultimately this method should be applicable not only to the identification of gene regulatory interactions, but to more complex tasks based on gene-gene correlations—such as the identification of cellular trajectories [40] and “tipping points” [41]—as well as providing a means to achieve a more principled approach to basic, early steps in scRNAseq analysis—such as normalization, batch correction, feature selection and clustering.

## Results

### The significance of gene-gene correlations

The statistical significance of correlations is rarely discussed because, for many common kinds of data—those that are continuous and at least approximately normal in distribution—the magnitude of correlation and its significance are related in a simple way that depends only on the number of measurements, and not the data distributions. Owing to Fisher [42], for any Pearson correlation coefficient (PCC), the *p*-value (probability of observing |PCC|≥*x* by chance) may be estimated as 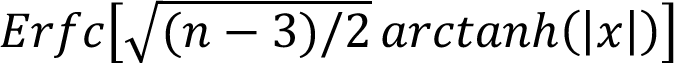, where *n* is the number of samples, and *Erfc* is the complement of the Error function (we refer to this expression henceforth as the “Fisher formula”).

Because scRNAseq data are both discrete and generally not normally distributed, *p*-values obtained using the Fisher formula cannot be accepted as accurate, but just how far off will they be? Figure 1 explores that question through simulation. Assuming Poisson-distributed data, and gene expression vectors of 500 cells in length, the formula does quite poorly for vectors of mean < 1, i.e., where the expected proportion of zeros exceeds 37%, assigning *p*-values that are too low for positive correlations, and too high for negative ones (Fig. 1A). For distributions with mean ≥1 (fewer than 37% zeros on average), the formula does reasonably well down to *p*-values as low as 10^-4^ but deviates progressively thereafter. This degree of accuracy would be a problem for any genome-wide analysis of correlations: To analyze pair-wise correlations among *m* genes one must test *m*(*m*-1)/2 hypotheses. With values of *m* often ≥ 12,000, this amounts to >7 x 10^8^ simultaneous tests, such that statistical significance of any single observation could potentially require *p* as low as 10^-9^, a value for which we may estimate, by extrapolation, that the Fisher formula is highly inaccurate even for genes with mean expression = 1.

**Figure 1.**
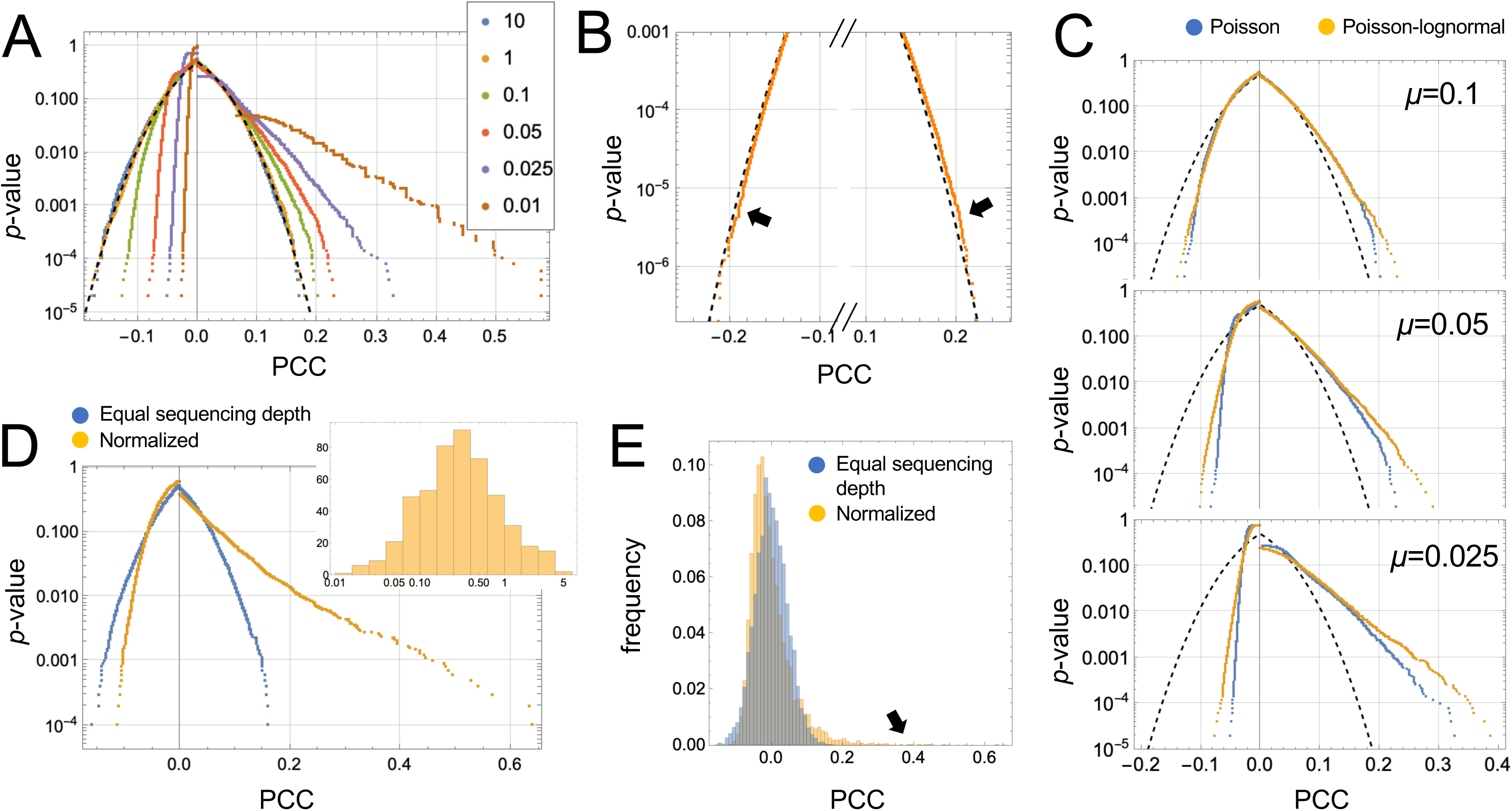
Relationship between Pearson correlation coefficient, vector sparsity and *p*-value. The panels compare *p*-values determined empirically by correlating 50,000 pairs of random, independent vectors of length 500 with *p*-values predicted by the Fisher formula. **A.** Data were independent random variates from Poisson distributions with means as indicated. The dashed line shows the output of the Fisher formula. **B.** Even for vectors drawn from a distribution with mean = 1, the Fisher formula significantly mis-estimated p-values smaller than 10^-4^. **C.** Poor performance of the Fisher formula is worsened when data are drawn from a Poisson-log-normal distribution, rather than a Poisson distribution (in this case the underlying log-normal distribution had a coefficient of variation of 0.5). **D-E.** Data were simulated under the scenario that gene expression is the same in every cell, but due to differences in sequencing depth, observed gene expression varies according to the depths shown in the inset to panel D. In panel D, true gene expression was adjusted so that observed gene expression after normalization would have a mean of 1, and the Pearson correlation coefficients (PCCs) obtained by correlating randomly chosen vectors are shown. Panel D plots empirically derived *p*-values as a function of PCC, whereas panel E displays histograms of PCCs. Compared with data that do not require normalization, associated *p*-values from normalized data are even more removed from the predictions of the Fisher formula.

The simulations in Fig 1A-B assume Poisson-distributed data but, as is often pointed out, scRNAseq data are usually over-dispersed relative to the Poisson distribution (more on this below). As shown in Fig. 1C, adding in such additional variance causes simulated data to deviate even further from the Fisher formula.

Further problems arise when considering that scRNAseq data always come from collections of cells with widely varying total numbers of UMI. Depending on the platform, such “sequencing depth” can vary over orders of magnitude, which is why normalization is usually considered a necessary early step in data analysis. Without normalization, it is obvious that many spurious gene-gene correlations would be detected, as any difference in sequencing depth between cells would, if not corrected for, induce positive correlation across all expressed genes.

As one might expect, normalizing individual reads by scaling them to each cell’s sequencing depth eliminates this bias, restoring the expected value of PCC under the null hypothesis to zero. Yet normalization does not restore the *distribution* of PCCs to what it would have been had all cells been sequenced equally. The consequences can be dramatic, as shown in Fig. 1D-E, where we simulate a case in which “true” gene expression is the same in each of 500 cells, but observed gene expression is a Poisson random variate from a mean that was scaled by a factor chosen from a distribution of cell-specific sequencing depths similar to what one might observe in a typical scRNAseq experiment, using the 10X Chromium platform (inset, Fig. 1D). Despite mean gene expression being ∼1, the relationship between PCC and *p*-value more closely resembles the case (in the absence of sequencing depth variation) where the mean is 0.01 (Fig. 1A). This is surprising, given that the average fraction of zeros in the normalized vectors was only 0.68 (which, for Poisson-distributed data, would be expected to occur at a mean expression level of 0.39). In short, for gene expression data resembling what is typically obtained in scRNAseq, the Fisher formula is highly unsuitable, for most pairs of genes, for estimating the significance of correlations.

### An analytical model for the distribution of correlations

The data in Figure 1 indicate that the relationship between PCCs and *p*-values is highly sensitive both to data structure and procedures intended to “correct” for technical variation. Because of this, we were concerned that a suitable null model for the distribution of correlations might be difficult to estimate empirically, especially when the true biological variation in most datasets is unknown. We therefore turned to constructing a null model analytically, attempting to account for known sources of variation (beside meaningful biological variation). The three sources considered were (1) variation introduced by imperfect normalization; (2) technical variation due to random sampling of transcripts during library preparation and sequencing; and (3) variation due to stochasticity of gene expression. The last of these cannot properly be called technical variation—fluctuating gene expression is a biological phenomenon—but like technical noise, gene expression fluctuation is usually an unwanted source of variation, and its effects need somehow to be suppressed.

With regard to the first source, we follow [43] in correcting not the gene expression data points themselves, but rather their Pearson residuals. Traditionally, the Pearson residual *𝑃_ij_* for cell *i* and gene *j*, is defined as

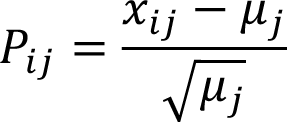

where *𝑥_ij_* is the gene expression value for cell *i* and gene *j*, and 𝜇_*j*_ is the mean expression of gene *j* averaged over all cells. As the Pearson residual is mean-centered, its expectation value, *𝐸*[*𝑃_ij_*], is zero; thus, the average of Pearson residuals for a large number of *𝑥_ij_* drawn from a single distribution should approach zero. The average of squares of Pearson residuals can be seen, by inspection, to approach the variance divided by the mean of the distribution from which the *𝑥_ij_* derive. Variance divided by mean is also known as the Fano factor and is often used to assess whether data are consistent with a Poisson distribution since, for any Poisson distribution regardless of mean, 𝐸[*P_ij_*^2^] = 1.

Pearson residuals may also be used to construct PCCs, which are commonly defined in terms of variances and covariances, but may be equivalently expressed as:

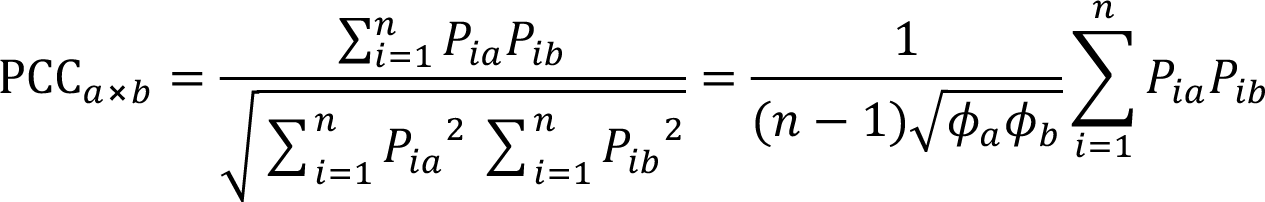

where PCC_$⨯&_ is the Pearson correlation coefficient between genes *a* and *b*, *n* the number of cells, and 𝜙_$_and 𝜙_&_ represent the Fano factors for genes *a* and *b*, respectively, i.e.

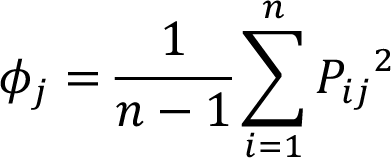

Since 𝐸[*𝑃_ij_*] = 0 for any gene, it follows that 𝐸[PCC_*axb*_] = 0, as long as all the expression values for gene *a* are drawn from a single distribution, and those for *b* are independently drawn from a single distribution.

However, in scRNAseq, unequal sequencing depth means that the expression values for any given gene are generally *not* drawn from a common distribution, but rather from one that is different for each cell. Interestingly, dividing *𝑥_ij_* in a Pearson residual by an appropriate scaling constant—i.e. normalizing the data—will restore 𝐸[*𝑥_ij_*] to 0, but will not restore the higher moments of *𝑥_ij_*, e.g., 𝐸[*𝑥_ij_*^#^] ≠ 1. To capture the correct second moment, one must scale the value of 𝜇_*j*_ inside each Pearson residual, rather than scaling *𝑥_ij_*. As [43] have pointed out, we can define a separate 𝜇_*ij*_ for each cell and gene:

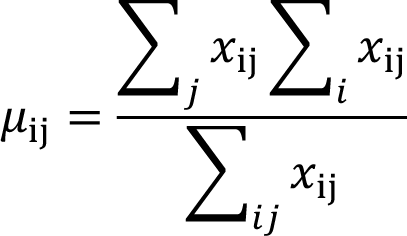

and consequently, define a corrected Pearson residual as

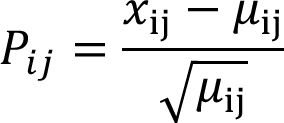

Although this transformation does not recover moments of the Pearson residual beyond the first two, it provides a principled alternative to traditional normalization. Moreover, by permitting calculation of a corrected Fano factor that has the appropriate expectation value under the assumption of Poisson distributed data, it can be used to test that assumption, in real data.

Source #2 refers to technical variation due to random, independent sampling of discrete numbers of transcripts. Although the sampling process in scRNAseq involves several discrete steps, including cell lysis, library preparation and DNA sequencing, several groups have argued that, at least at modest to low expression levels, simple Poisson “noise” can reasonably model the variation derived from these processes [35–38]. We accept this assumption here, but note that, in the following derivations, the Poisson distribution could just as easily be replaced by another known distribution, if it were adequately justified.

Finally, source #3 refers to the fact that “equivalent” cells are usually only equivalent in a time-averaged sense, i.e., transcript numbers will fluctuate around some mean value. Both theory and observation support the conclusion that these fluctuations can be large [30, 44–46]. The actual magnitude seems to differ for different categories of genes, but data from single-molecule transcript counting [44] suggest that, for most genes, the distribution of transcripts typically is approximately log-normal (consistent with the theoretical work of [47]), with a coefficient of variation in the range of 0.2-0.6.

Thus, even if library synthesis and sequencing performed identically across cells, one should not expect to observe Poisson-distributed reads. Under the reasonable assumption that gene expression fluctuations and sampling are independent, the variance of the combined process should be the sum of the variances of the composing processes. Since both processes have the same mean, we can re-state this thusly: the Fano factor of the combined process should be the sum of the Fano factors of the composing processes. One can then use this fact to further adjust the Pearson residual, and subsequently the Fano factor. Specifically, we define a “modified corrected Pearson residual” as:

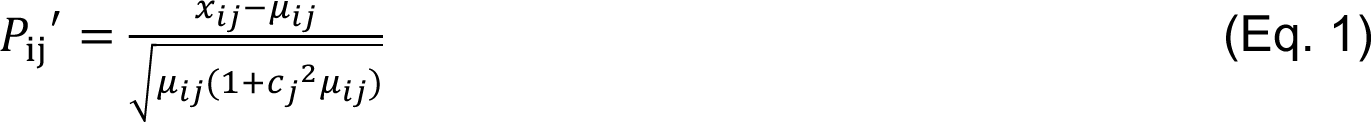

where *c* represents the coefficient of variation of gene expression for gene *j*. Accordingly, the expectation value for (*P*_ij_‣)^2^ becomes

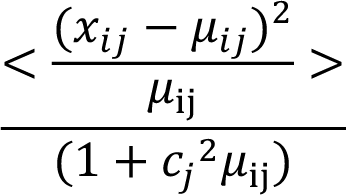

which resembles a Fano factor divided by (1 + 𝑐_*j*_^2^𝜇_ij_). Since 𝑐^2^𝜇_ij_ can alternatively be written as 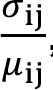, where *α* is the standard deviation of gene expression fluctuations, the term (1 + 𝑐_*j*_^2^𝜇_ij_) is simply the sum of the Fano factors for Poisson sampling (unity) and gene expression fluctuation (𝑐_*j*_^2^ 𝜇_ij_). Dividing by that sum essentially removes the additional variance due to gene expression noise from the expectation value of (𝑃_ij_,)^#^, restoring that value to 1.

In this way, one may define a “modified corrected Fano factor” 𝜙,equal to the expectation value of (𝑃_ij_,)^#^ . Likewise, we may use 𝜙, to define a modified corrected Pearson correlation coefficient, PCC′:

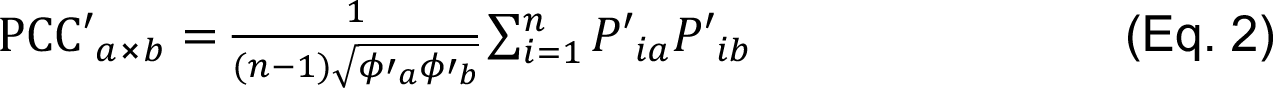

Notice that 𝜙, provides a measure of the degree to which a gene’s expression is more variable than expected by chance, and PCC′ provides a metric by which gene-pairs are judged more positively or negatively correlated than expected by chance, correcting in both cases for unequal sequencing across cells (without normalizing data) and the expected noisiness of gene expression.

To use these statistics in practice, one needs to know not only their expectation values but their full distributions under the null hypothesis. Constructing those analytically requires not only the coefficient of variation of gene expression noise, *c*, but the full distribution of that noise, which in the absence of information to the contrary, we will take to be log-normal [47], noting that any other distribution could as easily be substituted in the following discussion.

We thus treat *𝑥_ij_* as a Poisson random variable from a distribution whose mean is a log-normal random variable with a coefficient of variation of *c* (we refer to this compound distribution as Poisson-log-normal). Although an analytical form for the probability mass function of the Poisson-log-normal distribution is not known, we may derive analytical forms for an arbitrary number of its moments, as a function of *µ* (the mean) and *c* (see Methods). Thus, one can calculate for every cell *i* and gene *j*, given the observed values of *µ*_ij_, the moments of the expected distributions of the 𝑃_ij_, under the null hypothesis, and subsequently those for the (𝑃_ij_,)^#^. From there one can calculate the moments of any number of products and sums of 𝑃_ij_, and (𝑃_ij_,)^#^, such that, eventually, the moments of 𝜙, and PCC′ under the null hypothesis are ultimately obtained (see Appendix). Given a finite number of moments, one can estimate the tails of the distributions of these statistics (see Methods), allowing one to calculate the probability of extreme values of 𝜙,and PCC′ arising by chance (*p*-values).

This method, which we refer to as BigSur (<u>B</u>asic Informatics and <u>G</u>ene <u>S</u>tatistics from <u>U</u>nnormalized <u>R</u>eads), provides an approach for discovering genes that are significantly variable across cells (based on 𝜙,), and gene pairs that are significantly positively or negatively correlated (based on PCC′), automatically accounting for the widely varying distributions of these statistics as a function of gene expression level and vector length (number of cells). The one free parameter in the method, *c*, is relatively constrained, as its average value (over all genes) can be estimated from a plot of 𝜙, versus mean expression (see below). In this manner, one can avoid the use of arbitrary thresholds or cutoffs in deciding which genes are significantly “highly” variable (e.g., for dimensionality reduction and cell classification) and which genes are significantly positively and negatively correlated (e.g., to discover gene expression modules and construct regulatory networks).

### Performance on simulated data

In Figure 2 we simulate gene expression data for 1000 genes and 999 cells, under the null model described above (i.e., complete independence), varying “true” mean expression widely and uniformly over the genes, such that the most highly expressed genes average 3467 transcripts/cell and the most lowly expressed 0.0351 per cell. “Observed” gene expression values are then obtained by randomly sampling from a Poisson log-normal distribution with *c*=0.5, in which the gene-specific mean is first scaled in each cell according to a pre-defined distribution of sequencing depth factors (chosen to mimic typical ranges of sequencing depth when using the 10X Chromium platform). The result is a set of gene expression vectors of length 999, with means varying between 0.001 and 231.

**Figure 2.**
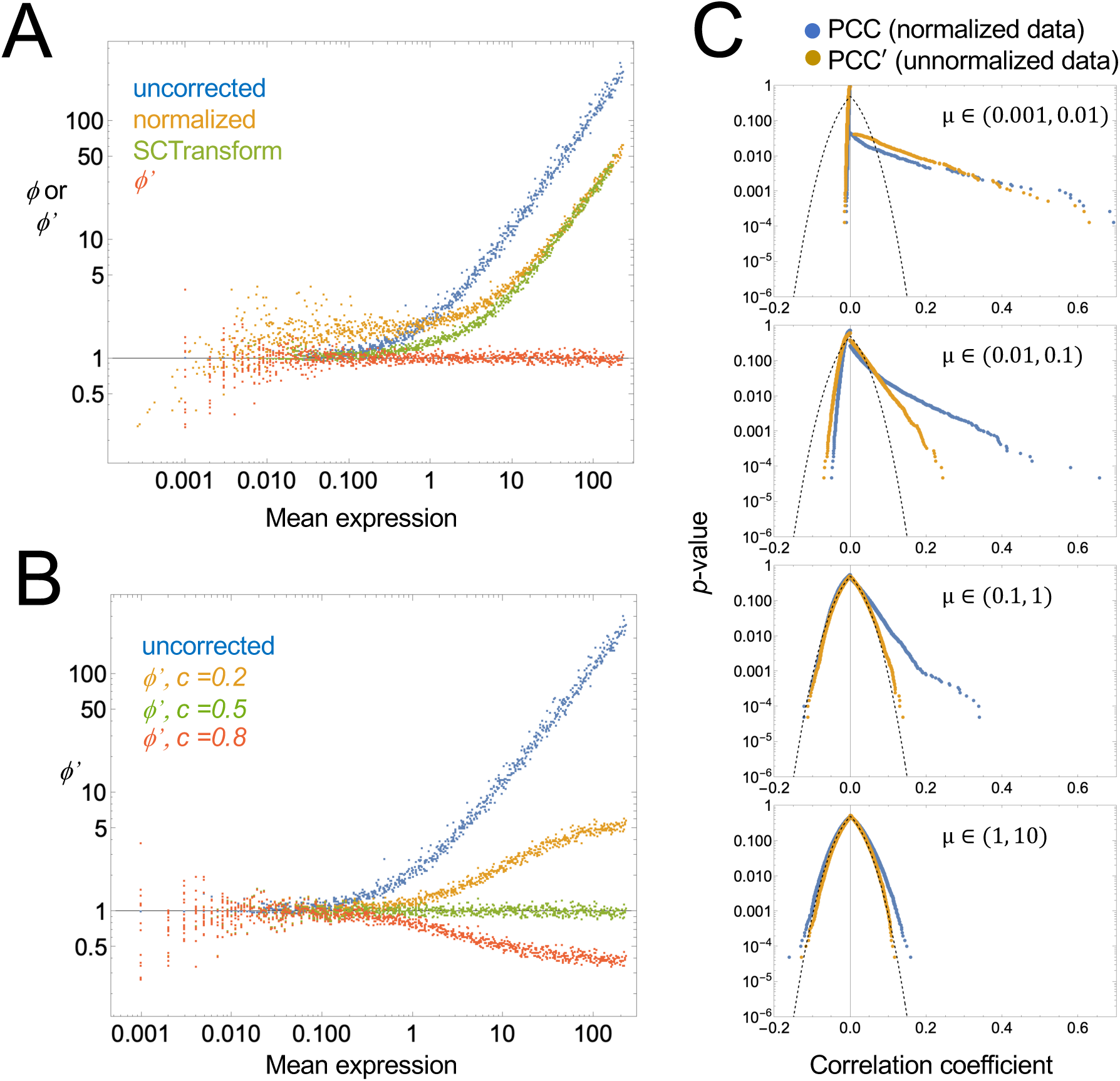
Comparing uncorrected and modified corrected Fano factors and correlation coefficients. Random, independent, uncorrelated gene expression data was generated for 1000 genes in 999 cells, under the assumption that observations are random Poisson variates from a per-cell expression level that is itself a random variate of a log-normal distribution, scaled by a sequencing depth factor that is different for each cell (see methods). **A.** Uncorrected (𝜙) or modified corrected (𝜙′) Fano factors are plotted as a function of mean expression level for each gene. Uncorrected factors were calculated either without normalization, or with default normalization (scaling observations by sequencing depth factors, learned by summing the gene expression in each cell). Uncorrected Fano factors were also calculated using SCTransform [48] as an alternative to default normalization. Modified corrected Fano factors were obtained by applying BigSur to unnormalized data, using a coefficient of variation parameter of *c*=0.5. **B.** Modified corrected Fano factors (𝜙′) were calculated as in A, but using different values of *c*. The data suggest that an optimal choice of *c* can usually be found by examining a plot of 𝜙,versus mean expression. **C.** Empirical *p*-values associated with uncorrected (PCC) or modified corrected (PCC′) Pearson correlation coefficients were calculated for pairwise combinations of genes in bins of different mean gene expression level (*µ*); examples are shown for four representative bins (both genes derived from the same bin). With increasing gene expression levels, the *p*-value vs. PCC relationship begins to approach the Fisher formula (dashed curve), but it does so much sooner for PCC′ than PCC.

As shown in Figure 2A, for most genes with mean expression greater than ∼0.1 reads/cell, uncorrected Fano factors exceed 1, and rise linearly with expression level. Normalizing the data—scaling expression in each cell to the relative sequencing depth of that cell— reduces high-expression skewing somewhat, but also elevates the Fano factors for genes with low expression, driving them closer to 2. These results, in which the majority of genes display Fano factors greater than 1, which rise further for highly expressed genes, agree with the pattern most commonly seen in actual scRNAseq data. Values of the Fano factor for lowly expressed genes may be restored to near 1 by normalizing using SCTransform, an algorithm designed to correct for some of the variance-inflating aspects of normalization-by-scaling [48], but the presence of high Fano factors among the highly expressed genes persists.

In contrast, if we calculate the modified corrected Fano factor, 𝜙′, for each gene, using *c*=0.5, we see that values are now centered around 1 at all expression levels (Fig. 2A). Note that choosing different values of *c* produces consistent positive or negative skewing at mean gene expression values above 1 (Fig. 2B). Under the assumption that most genes in real data should not vary significantly across cells, one may therefore estimate the optimal choice of *c* for any data set by simply finding the value that minimizes such high-expression skewing of 𝜙′.

Because it uses the moments of the Pearson residuals to calculate *p*-values, BigSur assigns statistical significance to every gene’s 𝜙′. As expected, given that the data in Fig. 2A were random samples, no values of 𝜙′ were found to be statistically significant at a Bonferroni-corrected *p*-value threshold of 0.05, or a Benjamini-Hochberg [49] false discovery rate of 0.05. Indeed, the lowest uncorrected *p*-value associated with any of the 1000 genes in Fig. 2A was 0.001.

Similarly, when the same synthetic data are analyzed for gene-gene correlations, one may compare the PCC values produced directly from normalized expression data with the PCC ′ values produced (from unnormalized data) by BigSur. As expected, both procedures return a distribution of values with zero mean, but PCC values are more broadly distributed than PCC′. Fig. 2C shows the frequency at which various values of the correlation coefficient arise when vectors with different mean gene expression are correlated. Although significant skewing from the Fisher formula is apparent, especially at low values of gene expression, it is much greater for PCC than PCC′. Indeed, for moderately expressed genes (e.g., 1-10 unique molecular identifiers [UMI] per cell), only PCC′ returns values whose distribution is relatively insensitive to expression level.

### Performance on real data

To characterize the performance of BigSur on real data, we used the droplet-based sequencing data of Torre et al., 2018 [50], obtained from a clonal isolate of a human melanoma cell line grown in culture. In this data set, the number of cells is large (8,640), data were validated on a second sequencing platform as well as by single molecule FISH, and the broad distribution of sequencing depths was typical of droplet-based sequencing.

We deliberately chose a clonal cell line because tissues always contain multiple cell types, i.e., groups of cells that express large sets of genes in cell-type specific ways. In such heterogeneous samples, genes that are associated with cell type identity will necessarily be strongly and densely correlated with each other; making the identification of correlations, in a sense, too easy—i.e., not a particularly good test of a method’s performance—and not particularly informative (one may expect to identify as correlated more or less the same genes one would find by clustering cells by gene expression and testing for differential expression between clusters).

In contrast, the use of (ostensibly) homogeneous cells forces BigSur to operate on more subtle connections—for example, those involving fluctuations in function-specific gene regulatory networks—that cell clustering and differential expression would not easily detect.

Accordingly, scRNAseq data from these cells were subjected to minimal pre-processing prior to analysis (see Methods), such that expression values were analyzed for 14,933 genes. Total numbers of UMI per cell varied greatly, ranging from 67 to 90,494, with 90% of cells containing between 666 and 9004 UMI.

First, we compared (uncorrected) PCCs for all gene-gene pairs, calculated using default-normalized expression data (i.e., data scaled to total UMI per cell), with PCC′ values returned by BigSur (Figure 3A-B). In panel B, frequencies are scaled logarithmically, to better display the distribution of large absolute values. BigSur associated a false discovery rate (FDR) of 2% to *p*-values less than 1.15 x 10^-4^, at which threshold it detected 639,789 correlations, 350,466 of which were positive. For uncorrected PCCs, the same *p*-value cutoff would translate, using the Fisher formula, to |PCC| > 0.041, which is displayed as a dotted line in Fig. 3B. Using this threshold, 1,484,156 correlations would be considered significant. Comparison of histogram shapes shows that use of uncorrected PCCs particularly inflates positive correlations, especially large ones, and under-counts negative ones, which is consistent with the results obtained using simulated data (Fig. 2C).

**Figure 3.**
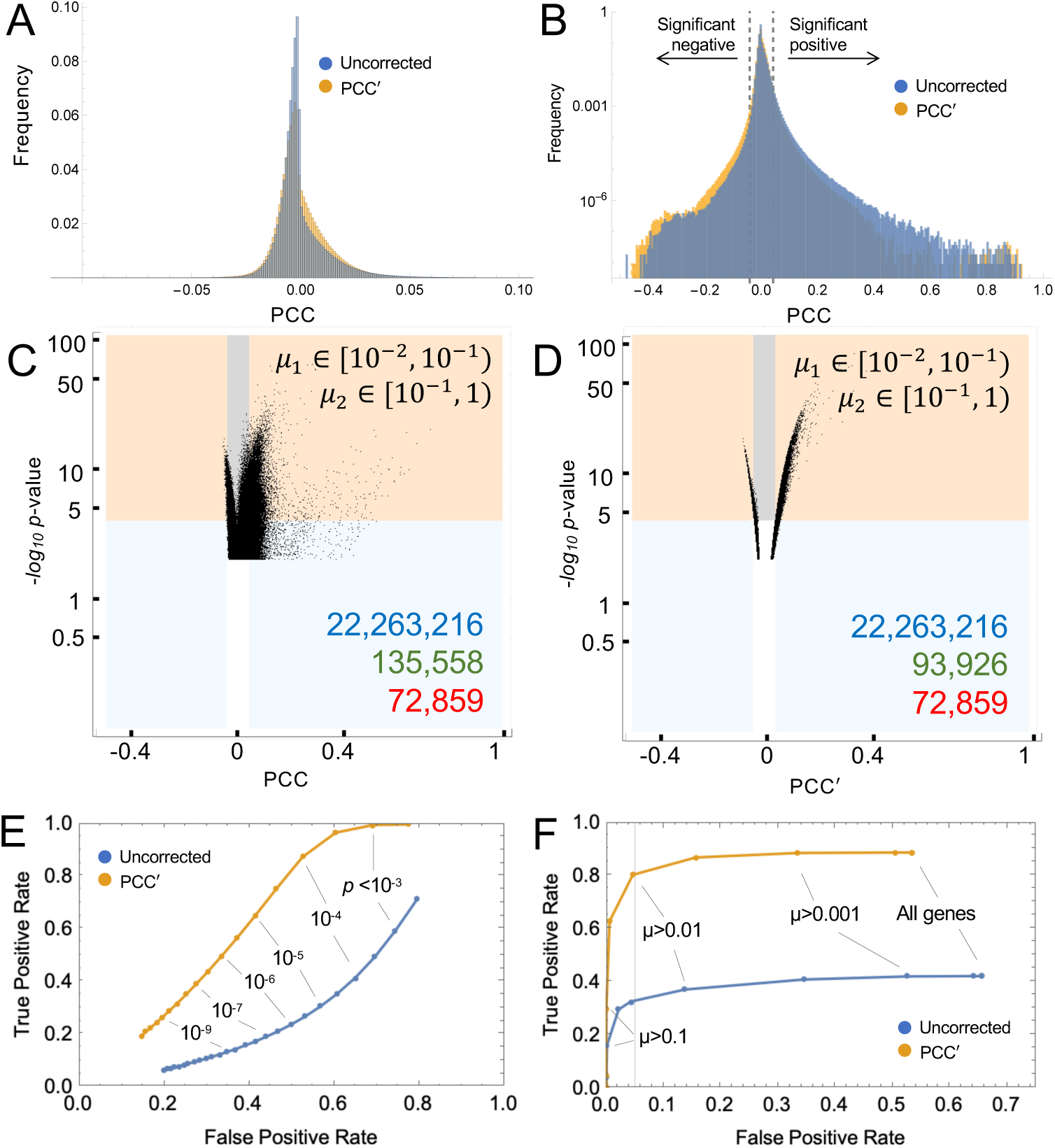
Statistical significance of pairwise gene correlations in data from a clonal cell line. A-B. Using scRNAseq data from a human melanoma cell line [50] (8640 cells x 14933 genes), pairwise values of PCC were calculated from normalized data, and PCC′ from raw data. Histograms display the frequency of observed values (the logarithmic axis in B emphasizes low-frequency events). Notice in B how positive skewing, also seen in simulated data (Fig. 2), is less for PCC′ than PCC. Dashed lines in B show thresholds at which Fisher formula-derived *p*-values would fall below 1.1 x 10^-4^. **C-D.** Scatterplots showing *p*-values assigned by BigSur to pairs of genes within two representative sets of bins of gene expression (for all pairwise combinations see Fig. S1-S2). The abscissa shows PCC (panel C) and PCC′ (panel D). The ordinate gives the negative *log_10_* of *p*-values determined by BigSur, i.e., larger values mean greater statistical significance. Orange and gray shading indicate gene pairs judged significant by BigSur (FDR<0.02). Blue and orange show gene pairs that would have been judged statistically significant by applying the Fisher formula to the PCC or PCC′, using the same *p*-value threshold as used by BigSur. The blue region contains gene pairs judged significant by the Fisher formula only, while the unshaded region shows gene pairs not significant by either method. Numbers in the lower right corner are the total numbers of possible correlations (blue), statistically significant correlations according to the Fisher formula (green), and statistically significant correlations according to BigSur (red). **E-F.** ROC curves assessing whether the overall performance of the Fisher formula—applied either to PCC or PCC′— can be adequately improved either by using a more stringent *p*-value cutoff (E) or limiting pairwise gene-correlations to those involving only genes with mean expression above a threshold level, *µ* (F).

To see how the discovery of correlations varied with gene expression level, we divided genes into 6 bins of different mean expression, and separately analyzed correlations between genes in all 21 possible combinations of bins. The full data are presented in Fig. S1-S2, with two representative panels in Fig. 3C-D. Each point represents a gene-gene pair. The value on the abscissa is either the uncorrected PCC (Fig. 3C and S1) or modified corrected PCC′ (Fig. 3D and S2), and the value on the ordinate is the -*log_10_ p*-value, as determined by BigSur (i.e., the larger the number, the lower the *p*-value). Shaded territories mark those data points that were judged statistically significant (FDR<0.02) either by Big Sur (orange and gray), or when *p*-values were calculated by the Fisher formula (blue and orange; for further details see figure legend). The data confirm that the Fisher formula returns an excess of correlations, compared to Big Sur, albeit less severely for PCC′ than for PCC. Examination of the full dataset (Fig. S1-S2) shows that many more truly significant correlations are found among highly expressed genes; and the Fisher formula performs worst when at least one of the pairs in a correlation is a lowly-expressed gene. Indeed, as gene expression becomes high (e.g., mean value >1 UMI per cell for both genes in a pair), the distribution of *p*-values calculated by BigSur for PCC′ begins to approximate the Fisher formula reasonably well (Fig. S2), with deviation apparent only for very low *p*-values (-*log_10_p* > 10). This observation validates the accuracy of the method used by BigSur to recover *p*-value distributions from the first five moments of the expected distributions of modified, corrected Pearson residuals (see Methods).

Assuming, for the sake of illustration, that the correlations judged significant by BigSur represent ground truth, we may then calculate levels of true- and false-positivity when *p*-values are calculated by feeding either PCC or PCC′ into the Fisher formula. This enabled us to ask whether applying a more stringent *p*-value cutoff, or thresholding gene expression (i.e., excluding genes below a certain expression level), might enable this simpler, formulaic approach to achieve an acceptable level of sensitivity and specificity. As shown by the receiver-operating characteristic (ROC) curves in Fig. 3E, the performance of uncorrected PCCs is exceedingly poor regardless of *p*-value threshold, with false positives exceeding true positives at all values. PCC′ does better, but to control FDR to <10%, one still loses the ability to detect >85% of true positives.

Arbitrarily thresholding gene expression performs somewhat better (Fig. 3F). For uncorrected PCCs, one must exclude all genes with expression < 0.065 UMI/cell to control the FDR at 5%, which for this dataset means discarding 67% of all gene expression data, and in return recovering only 33% of true positives. PCC′ does much better here: we may recover 81% of true positives at an FDR of 5% by discarding the ∼52% of genes with the lowest expression. While the exact numbers are likely to vary for different data sets, these observations suggest that, if one is willing to sacrifice the power to identify a substantial minority of correlations, feeding PCC′ (but not PCC) into the Fisher formula can represent an acceptable and computationally fast alternative to *p*-value identification by BigSur.

### Clustering using correlations

Although BigSur found 639,789 statistically significant correlations in this dataset (about 0.57% of all possible pairwise correlations) the vast majority had quite small values of PCC′ (Fig. 3A-B), indicating that most statistically significant correlations were weak. To obtain a measure of correlation strength that could be compared across samples of different lengths (numbers of cells), we expressed each correlation in terms of an “equivalent” PCC, which is simply the PCC value that, for continuous, normally-distributed data of the same length, would have produced the same *p*-value (by the Fisher formula). As shown in Fig. S3, only 4335 gene pairs displayed equivalent PCCs greater than 0.2 or less than -0.2.

Yet, despite the weak strength of most correlations, there are good reasons to believe them to be biologically relevant. One of the simplest comes from examining the frequency at which correlations arise among paralogous genes and genes that encode proteins that physically interact. It is known that gene paralogs are often co-regulated [51] leading us to expect paralog pairs to be enriched among *bona fide* correlations. It is also reasonable to expect that transcripts encoding proteins that interact will be co-expressed at least some of the time. As it happens, among the correlations identified by BigSur we observe ∼12 fold enrichment for paralogs and ∼7.5 fold enrichment for genes encoding physically-interacting proteins [see Methods].

To divide the full set of correlations into potentially interpretable groups, we used a random-walk algorithm [52] to identify subnetworks more highly connected internally than to other genes; we refer to these as gene communities. Most communities were of modest size, with 94 of the 96 largest containing between 4 and 392 genes each. However, the largest two contained 2160 and 1570 genes respectively, were very densely connected internally, and strongly anti-correlated with each other (Figure 4A). These factors strongly suggest that these cells, despite being a clonal line, are heterogenous, falling into (at least) two distinct groups. Interestingly, the largest gene community contained virtually the entire set of mitochondrially-encoded mitochondrial genes, and the second largest contained virtually all protein-coding ribosomal genes (for both cytoplasmic and mitochondrial ribosomes). Using the top 50 most-highly connected genes (those with the largest positive equivalent PCCs) in each of these communities as features, we performed Leiden clustering on the modified corrected Pearson residuals of all 8,640 cells, and easily subdivided them into three clusters of 5,812, 1,180 and 1,638 cells, which we labeled as clusters 1,2 and 3, respectively (Fig. 4B; see Methods).

**Figure 4.**
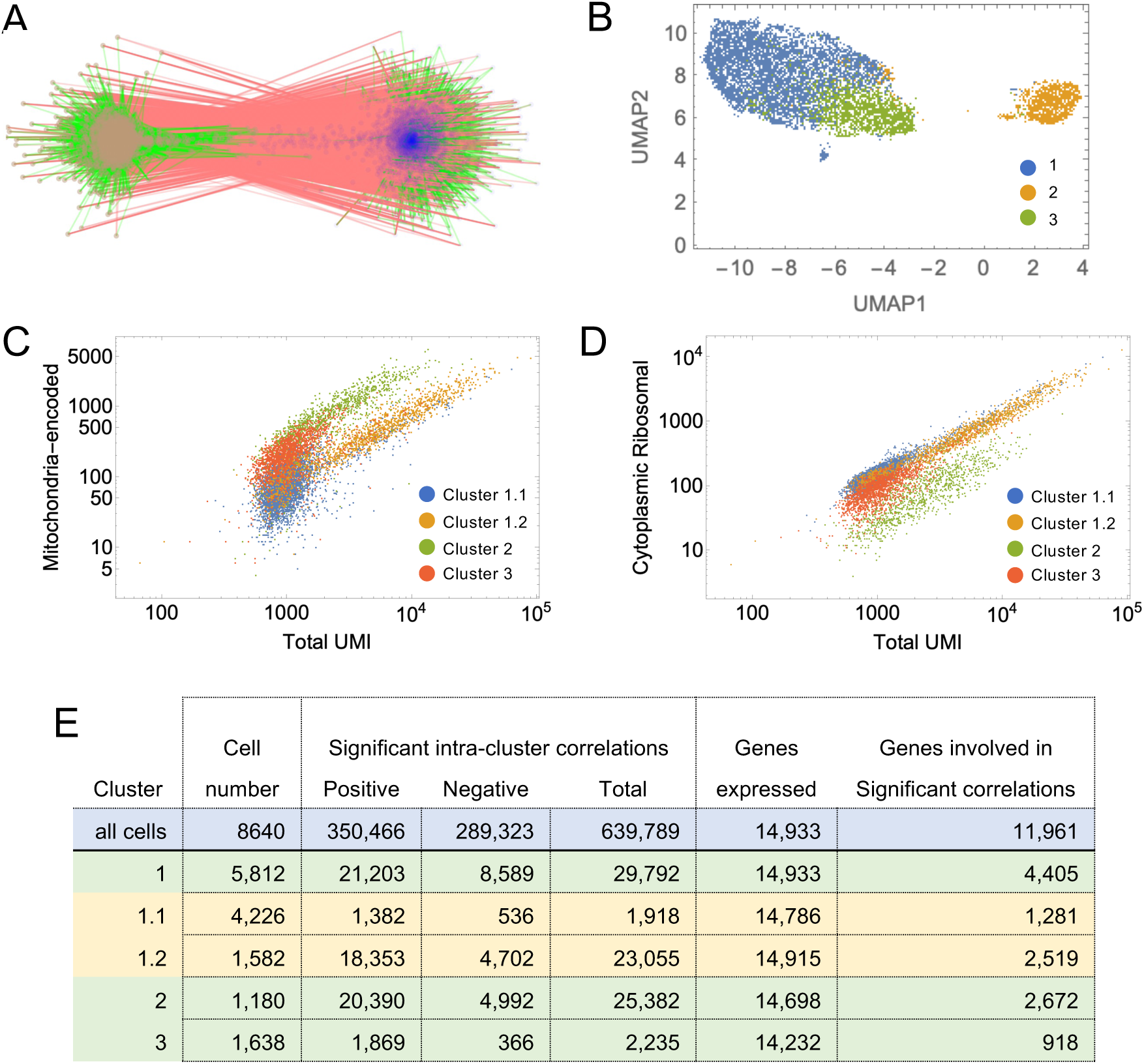
**Clustering cells based on mitochondrial and ribosomal communities. A**. Correlations among the two largest gene communities detected by BigSur are shown. Vertices are genes, and edges—green and red—represent significant positive and negative correlations, respectively. Blue vertices represent members of the mitochondrially encoded gene community and brown vertices the ribosomal protein gene community (Labels have been omitted due to the large numbers of gene involved). **B.** Using the top 50 most highly positively connected vertices in the two communities as features, cells were subjected to PCA and Leiden clustering; the three clusters recovered are displayed by UMAP. **C-D.** After a second round of clustering of Cluster 1, the resulting four cell groups were analyzed for the distribution of expression of ribosomal protein-coding and mitochondrially-encoded genes, as a function of total UMI in each cell. The results show that the four clusters form distinct groups based on their relative abundances of ribosomal and mitochondrial genes. **E.** Results of applying BigSur to each cluster. Note the large decrease in statistically significant correlations—especially negative correlations—in any of the clusters when compared with results obtained using all of the cells together (>600,000 total correlations). This is consistent with heterogeneity in the original sample, causing large blocks of genes to correlate and anti-correlate.

We then analyzed each cluster independently by BigSur, re-calculating PCC′ values and statistical significance for each group of cells. Surprisingly, in each cluster BigSur again found two large, strongly anti-correlating communities, one of which contained the mitochondrial-encoded genes and the other the ribosomal protein genes. This led us to further subcluster cluster 1, again using the 50 most-highly connected genes in each of these two communities as features and subdivided it into two groups of 4,226 (cluster 1.1) and 1,582 cells (cluster 1.2).

Variable expression of mitochondrially-encoded genes is a common finding in scRNAseq. Their presence at high levels (e.g. >25% of total UMI) is usually interpreted as an indication of a “low quality” cell—potentially one in which the plasma membrane has ruptured and cytoplasm has been lost—or perhaps a cell in the process of apoptosis [53]. Closer examination of the cell clusters identified in this dataset suggests these phenomena are likely only part of the explanation. In Fig 4C, we plot mitochondrial-encoded UMI versus total UMI for the entire set of cells, coloring them according to the four cell clusters mentioned above. Several distinct behaviors were noted. Cell cluster 2 forms a coherent group with high mitochondrial expression that is linearly proportional to total UMI. Throughout this group, the percentage of total transcripts that is mitochondrial remains in a narrow band, with mean of 34% and coefficient of variation (CV) of 0.41. Cluster 3 displays low total UMI, and a mitochondrial fraction centered around a mean of 21% (CV=0.43). Cluster 1.2 has high mitochondrial expression, but even higher total UMI, such that the mitochondrial fraction averages 9.7% (CV=0.45), while cluster 1.1 has both low total UMI and low mitochondrial UMI (average mitochondrial fraction 8.2%, CV=0.47).

In Fig. 4D, we also plot total UMI derived from ribosomal protein genes against total UMI. Of all the clusters, cluster 2 best fits the expectation for “damaged” cells: Mitochondrial and ribosomal UMI rise proportionately with total UMI (consistent with randomly varying sequencing depths), but the proportion of mitochondrial UMI is almost three times higher, the proportion of ribosomal UMI almost three times lower, and total UMI about 2.5 times lower than in cluster 1.2, the only other cluster that displays a wide range of sequencing depths. Yet it is curious that the mitochondrial proportions in cluster 2 are so narrowly distributed around a mean; one might expect variable degrees of cytoplasm leakage following cell damage to produce a broad distribution beginning near cluster 1.2 and gradually tapering off. The absence of such behavior suggests that such cells are not merely variably damaged during preparation, but represent a distinct, possibly pre-existing state, perhaps associated with some form of cell stress or death (although we see no significant enrichment of gene expression associated with apoptosis [54] in cluster 2).

Even if we remove cluster 2 from further analysis, clusters 1.1, 1.2 and 3 also differ between in each other in relative proportions of mitochondrially-encoded, ribosomal and total transcripts; however, the way in which they do so is not suggestive of any simple mechanism of cell damage (and overall mitochondrial content is not in the range that would necessarily lead to exclusion from downstream scRNAseq analyses). As mentioned above, using the genes in the mitochondrially-encoded- and ribosomal-enriched communities as features, analysis of gene correlations using BigSur separately on each of the four clusters revealed anti-correlating communities of mitochondrially-encoded and ribosomal genes in every case (Figure 5A-D; Table 2). In fact, even after another round of subclustering of the subclusters derived from cluster 1 (cluster 1.1 and cluster 1.2) using these genes as features, BigSur still identified distinct, anti-correlating mitochondrially-encoded and ribosomal communities (not shown). These data strongly suggest that, notwithstanding effects of cell damage or cell death, there exists among healthy cells continuous, anti-correlated variation in both the mitochondrially-encoded and ribosomal protein genes, suggestive of some biologically meaningful regulatory relationship (discussed further below).

**Figure 5.**
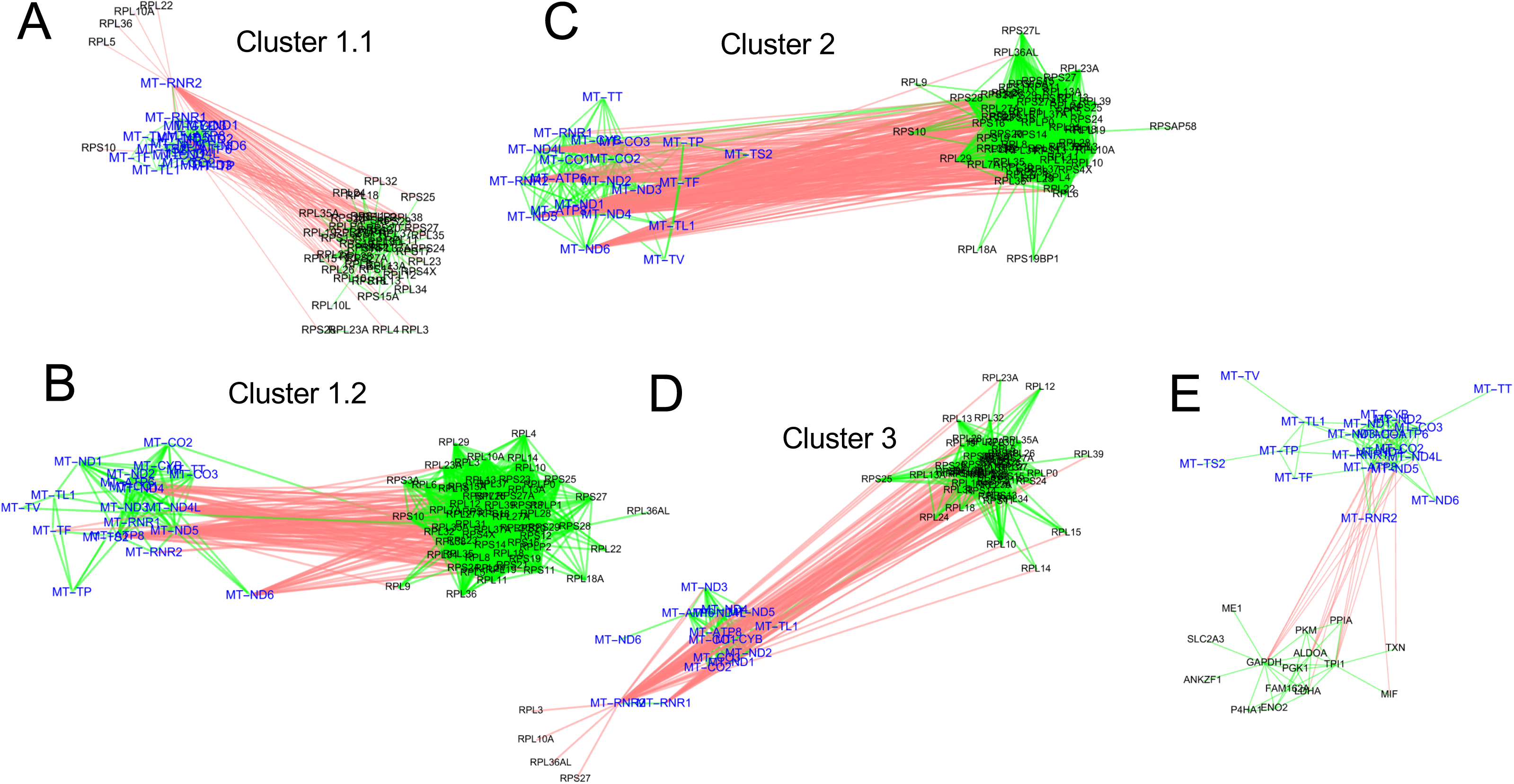
Mitochondrial communities are a source of strong anti-correlations. Mitochondrially-encoded and ribosomal protein gene communities were identified in all clusters. Anti-correlations identified specifically between mitochondrially-encoded genes and ribosomal protein genes within the different clusters are shown in A (cluster 1.1), B (cluster 1.2), C (cluster 2) and D (cluster 3). Panel E shows additional anticorrelations in cluster 1.2 involving mitochondrially-encoded genes and glycolysis genes For ease of readability, mitochondrial genes have been highlighted in blue. Red links refer to negative correlations; green to positive.

### Analysis of gene communities

Figure 4E summarizes the results of using BigSur to identify significant correlations (FDR<0.02) in each of the cell clusters described above (1.1, 1.2, 2 and 3). Because statistical power to detect correlations falls with number of cells analyzed, one might expect to see the fewest significant correlations in the smallest clusters, but this was not the case. Instead, the clusters with the largest number of significant correlations were those with the highest average sequencing depth (Fig. 4C), suggesting that data sparsity has an especially strong influence on correlation detection.

Schadt and colleagues have recently argued [55] that negative correlations may be considered a strong indicator of cell heterogeneity. These authors argue that minimization of the proportion of negative gene-gene correlations provides a principled metric for determining when to cease sub-clustering cells. Consistent with this view, we observed that, as cells were successively subclustered, the proportion of negative correlations fell from 45% to about 20% (Fig. 4E). The clusters with the lowest proportion of negative correlations were clusters 1.2, 2 and 3. As cluster 2 had very high levels of mitochondrially-encoded genes (33.7% of UMI), and cluster 3 had the lowest average UMI/cell (1,273) of any cluster, we focused our analysis on cluster 1.2, which had both low levels of mitochondrial genes (9.7% of UMI) and high average UMI/cell (6,252). .

Within cluster 1.2, we found that enrichment for correlated paralog pairs and genes whose products interact physically was much higher than when all 8,640 cells had been considered together. Specifically, among the positive correlations, paralog pairs were enriched 55-fold, and links supported by protein-protein interactions 32-fold, over what would have been expected by chance. Indeed, 15% of all positive correlations in cluster 1.2 corresponded to links supported by known protein-protein interactions.

Table 1 and Figures S4-S7 summarize results for 13 of the largest gene communities detected in cluster 1.2, which collectively account for 1,291 of the 2,519 genes that showed any significant positive correlations. Table 1 groups the genes of each community into functional categories according to annotations found in the most recent release of the MSigDB database [56, 57]. In 11 of the 13 communities (accounting for 1,217 of the 1,291 genes), over 50% of the genes could be associated with just one or a few functional annotations (Table 1).

**Table 1:**
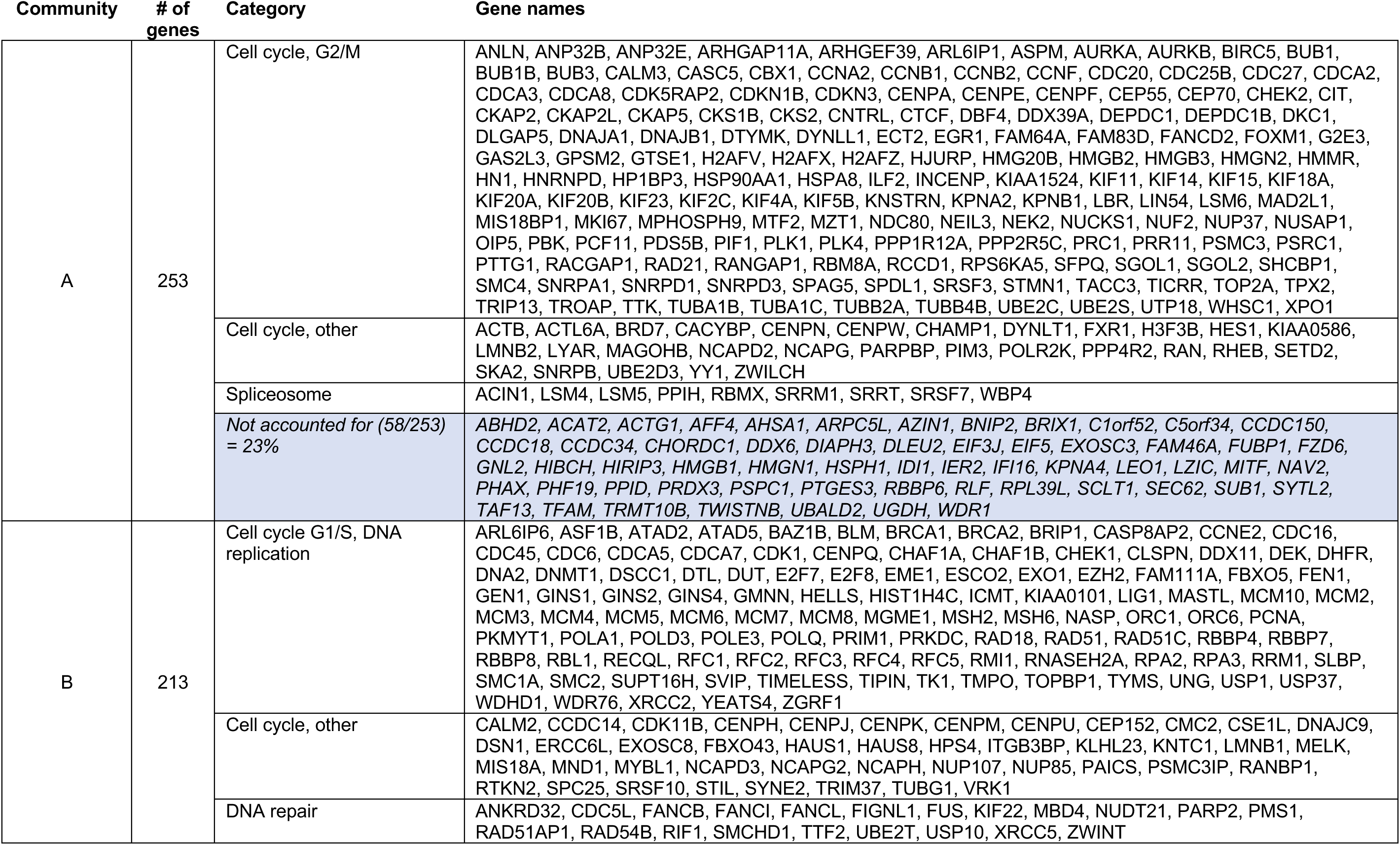
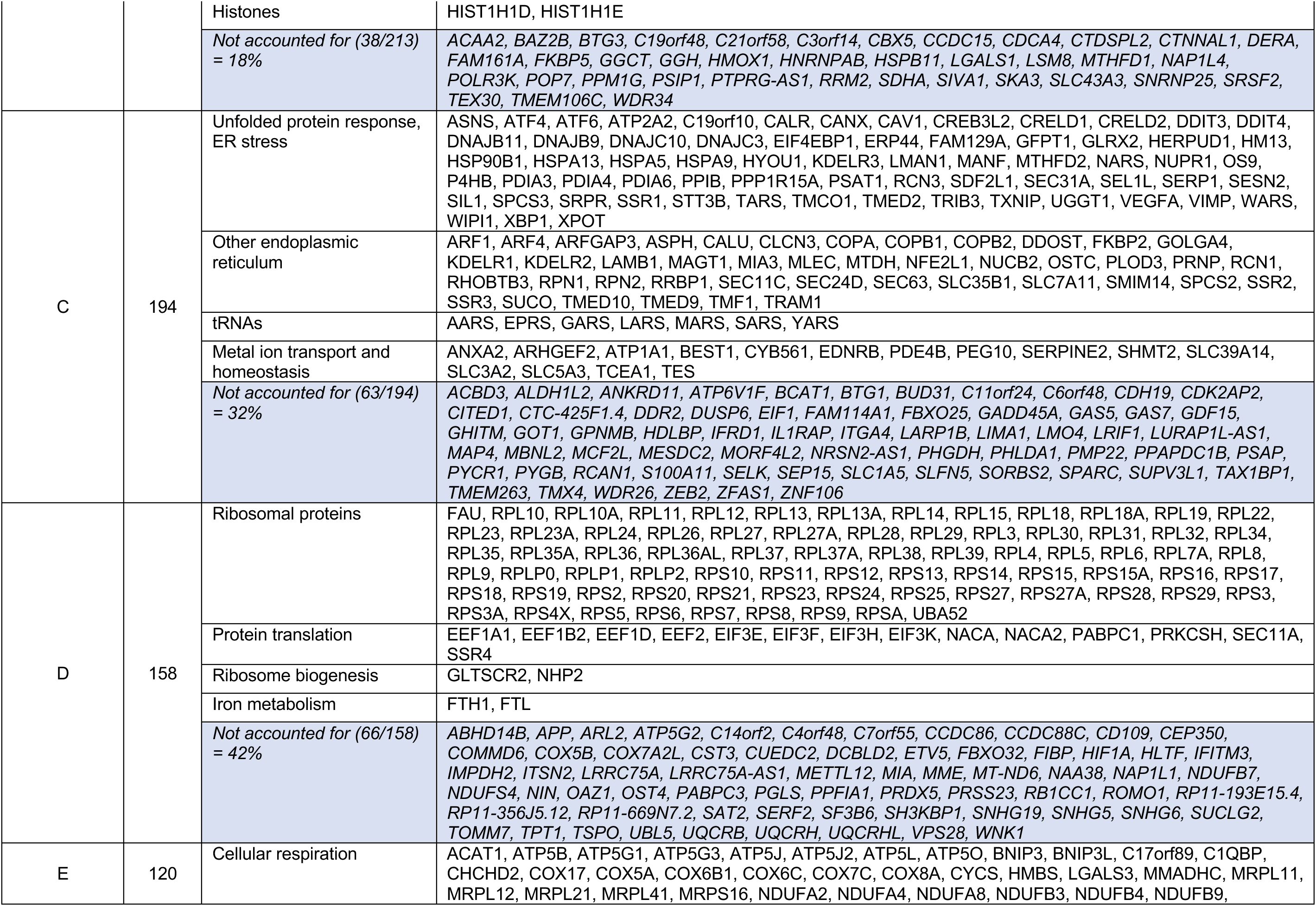
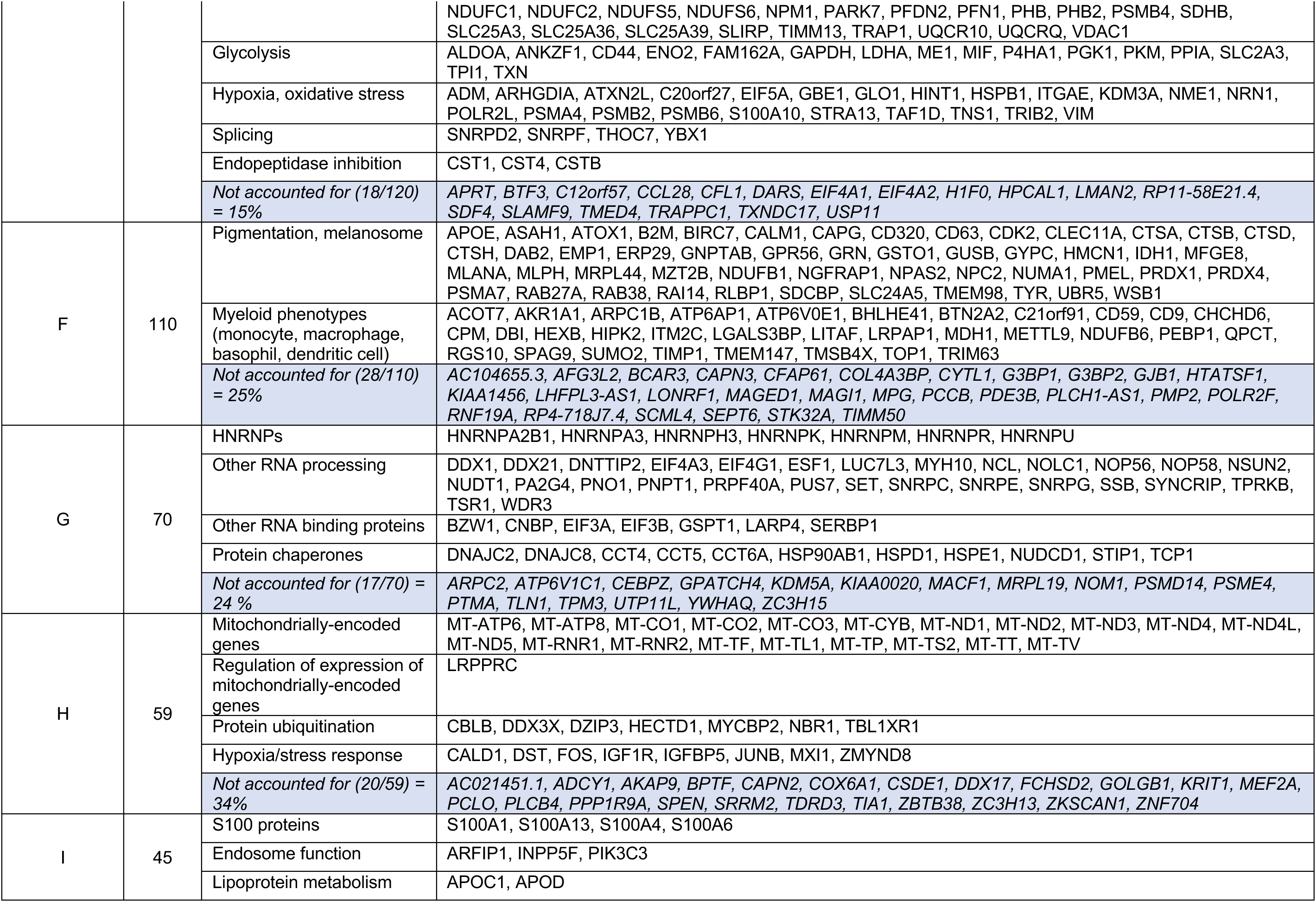
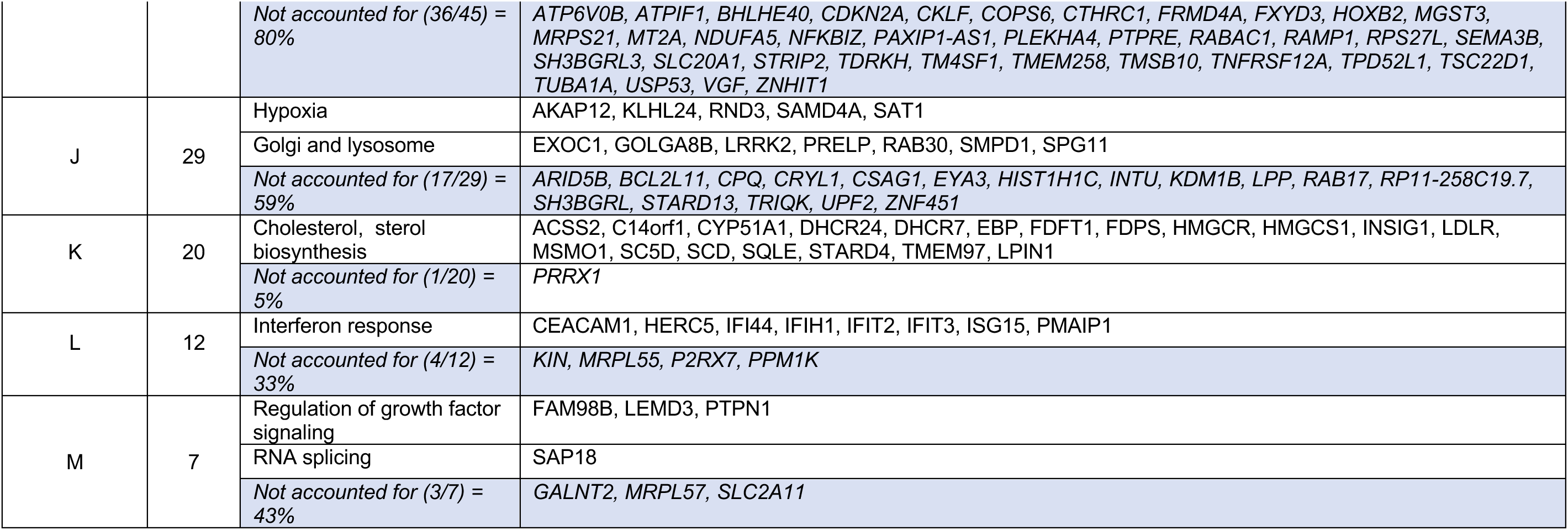
Gene communities identified using cell subcluster 1.2. Communities containing at least 7 genes are shown. Genes were assigned to annotations after manually exploring overlaps with all MSigDB datasets (category labels used here often reflect the merger of redundant or semi-redundant gene sets). Genes marked “Not accounted for” failed to overlap significantly with known gene sets (i.e., overlaps involved at most two genes, or accounted for less than 1% of genes in the dataset).

For example, communities A and B consist primarily of genes related to the cell cycle, with more than 62% of the genes in community A known to be preferentially associated with the G2 and M phases of the cell cycle. About 71% of community B is associated with the cell cycle, the majority of these being associated with G1 and S phase. These communities were also correlated with each other, but the community-finding algorithm easily subdivided them.

Most other communities are at best weakly correlated with either of the cell-cycle communities. In community C, 34% of genes encode proteins involved in the unfolded protein response and/or endoplasmic reticulum (ER) stress, and other ER proteins account for another 22% of genes. ER stress is commonly observed in cancer, and the unfolded protein response is specifically and strongly activated in melanoma cells [58–60]. Community D contains almost all genes encoding protein components of cytoplasmic ribosomes, plus a large set of genes that regulate ribosome biogenesis or function. Overall, ribosome-related genes make up 57% of this community. Community E combines genes involved in cellular respiration, glycolysis, and oxidative and hypoxic stress, which collectively account for 78% of this community.

Fifty genes in community F, nearly half the total, are associated with melanin synthesis and melanosome biogenesis, and include traditional melanocyte markers such as *MLANA*, *PMEL*, *RAB27A* and *TYR*. Another subset in community F, consisting of 32 genes, shares annotations related to markers of myeloid cell types (monocytes, macrophages, basophils, etc.) but largely consists of ubiquitously expressed genes involved in processes such as lipid biosynthesis. At least one of these genes, *TRIM63*, is known to be strongly associated with melanoma [61], and is a validated target of MITF [62], the primary transcription factor controlling expression of pigmentation genes.

Community G contains multiple genes involved in RNA processing, including seven genes encoding members of the ubiquitously expressed heterogeneous nuclear ribonucleoprotein (HNRNP) family, as well as genes encoding protein chaperones of the HSP10, HSP40 HSP60, HSP90, and CCT families. Community H contains most of the mitochondrially-encoded genes, as well as genes involved in protein ubiquitination and the hypoxia stress response.

Community I combines four genes encoding S100-family calcium binding proteins, as well as various genes associated with endosome function and lipoprotein metabolism. Unlike other communities, in this community most genes do not fall into large groups with traditional annotations. However, many of them are genes known to be strongly associated with melanoma. These include APOC1, APOD, *CDKN2A*, *CTHRC1*, *FXYD3*, *INPP5F*, *MT2A*, *S100A4, SEMA3B, TMSB10* and *VGF* [63–66], suggesting this community is detecting genes co-regulated by drivers of a melanoma-specific cell state.

Community J combines genes associated with the response to hypoxia with genes involved in Golgi body and lysosome function. Nearly all the genes in community K are associated with cholesterol/sterol biosynthesis and homeostasis; the only exception is *PRRX1*, which serves as the transcriptional co-regulator of serum response factor. In oligodendrocytes at least, it has been shown that PRRX1 is necessary for the expression of cholesterol biosynthesis genes [67].

Community L consists mainly of genes annotated as related to interferon signaling; these genes are also the primary drivers of the cellular anti-viral response. Finally, community M contains several genes associated with growth factor signaling.

Figures S4-S7 display the gene-gene linkages within these communities graphically. Here, genes are shown as light blue disks, except for transcription factors which appear as yellow squares. The areas of the disks and squares are proportional to the relative mean expression of the genes (absolute scaling differs between panels, as it was adjusted to enhance the readability of each figure). Green lines denote significant positive correlations (no negative correlations were observed in any of the communities shown). In several cases, a smaller version of each graph, in which links supported by protein-protein interaction data have been overlayed with brown lines, is displayed as an inset; for some communities, only the graphs containing these highlights are shown. Examination of Fig. S4-S7 shows that genes associated with a single functional annotation, or that encode directly-interacting proteins, are often especially densely interconnected, causing them to cluster together (e.g., genes encoding directly physically interacting proteins in A and B, unfolded protein response genes in C, ribosomal subunit genes in D, glycolysis genes in E, S100 genes in I, etc.).

Whereas the clustering of genes into communities had been carried out based on positive gene-gene correlations, plotting pairs of communities together enabled the evaluation of negative correlations between them, the vast majority of which involved the mitochondrially-encoded genes which, as mentioned previously, strongly anti-correlated with ribosomal protein genes (Fig. 5). Mitochondrial genes also anti-correlated with some of the genes in communities A, F, E and G; for example, anticorrelations between mitochondrial genes and genes encoding glycolytic enzymes are shown in Fig. 5E.

Figure 5 also documents that the anti-correlation between mitochondrial and ribosomal genes is as apparent in cell clusters 1.1, 2 and 3 as it is in cluster 1.2. This is interesting insofar as the features used to subdivide cells into these clusters included most of mitochondrially-encoded and ribosomal genes. What this suggests is that there are not two separable populations of cells expressing mitochondrial versus ribosomal genes. Instead, the data suggest that the observed anti-correlations are not themselves sufficiently correlated with each other to drive cell clustering. In other words, BigSur seems to able to detect local gene-gene relationships that could not have been identified by comparing differential expression of genes in any possible grouping of cells.

### Estimating the accuracy of BigSur

The association of a small number of annotation categories with nearly all of the gene communities that BigSur found, in an unsupervised manner, in scRNAseq data strongly suggest that many of the correlations detected by BigSur are true positives. Nevertheless, in nearly all communities some genes did not fall into obvious categories (Table 1). Do these just reflect the incompleteness of existing annotations, or did BigSur produce false positive results beyond the ∼2% expected from the FDR cutoff that was used?

Rarely in biology does one have ground truth information with which to settle this question, but one can still perform informative tests. For example, one can consider a subset of the genes that form a functional category and ask whether BigSur correctly identifies other members of the category as being correlated with them. An example is shown in Figure 6A, where we started with a 27-gene panel, the “Reactome Cholesterol Biosynthesis” gene set from MSigDB (*ACAT2, ARV1, CYP51A1, DHCR24, DHCR7, EBP, FDFT1, FDPS, GGPS1, HMGCR, HMGCS1, HSD17B7, IDI1, IDI2, LBR, LSS, MSMO1, MVD, MVK, NSDHL, PPAPDC2, PMVK, SC5D, SQLE, SREBF1, SREBF2, TM7SF2)*. That panel includes many, but not all, genes known to be involved in cholesterol metabolism. The graph displays all statistically significant positive correlations involving any of these genes that were detected in cell cluster 1.2 (members of the gene panel are indicated with large font blue lettering).

**Figure 6.**
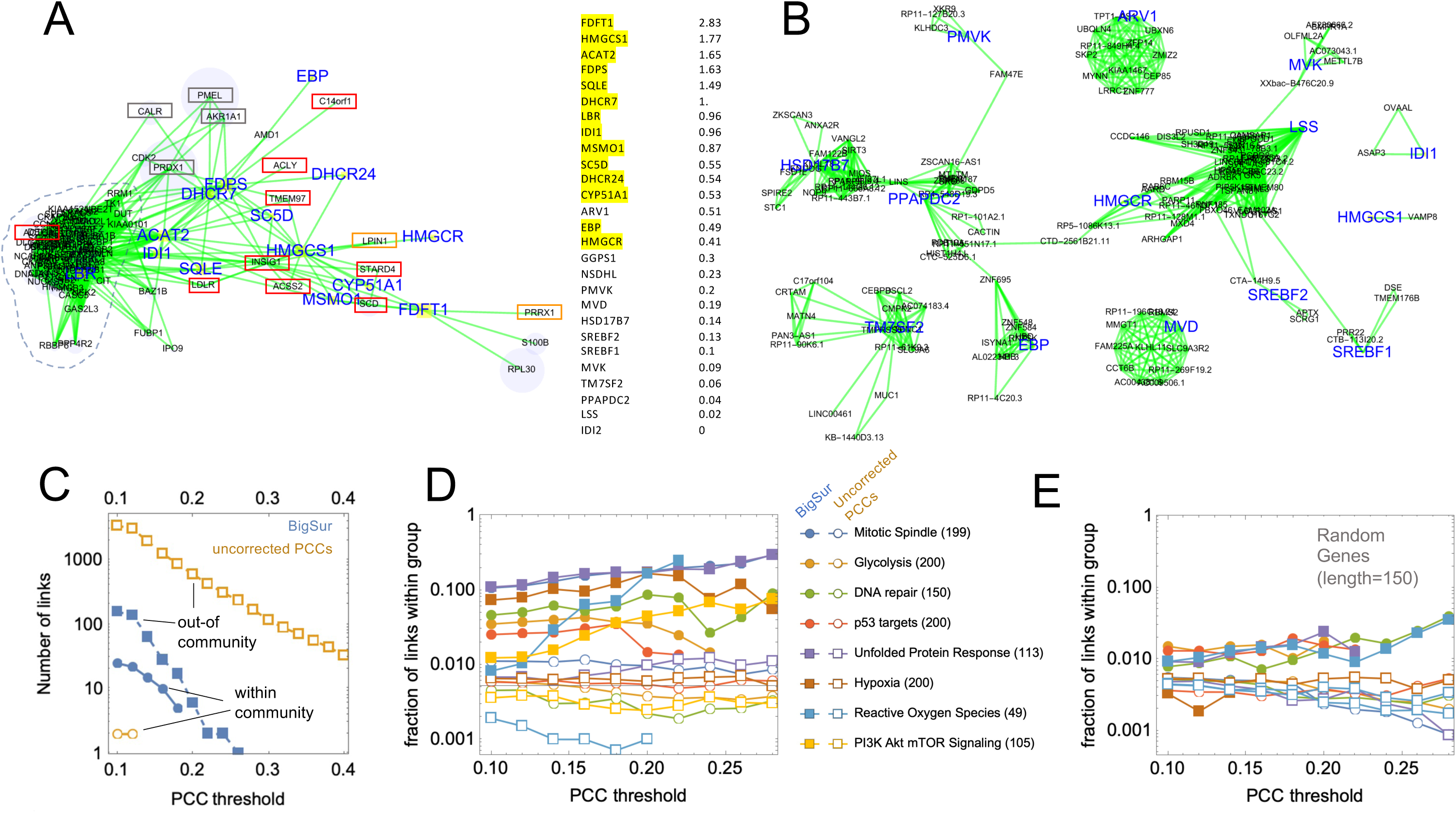
Comparing the ability of BigSur and uncorrected PCCs to identify biologically significant correlations. Using the data from cell cluster 1.2, BigSur identified positive correlations and their *p*-values for all genes, converting the *p*-values to equivalent PCCs. In addition, the same starting data were also normalized and uncorrected PCCs obtained. A-D compare the correlations identified by the two methods. **A.** Genes identified by BigSur as correlating with the Reactome Cholesterol Metabolism gene set. Genes belonging to the set are highlighted with boxes; additional known cholesterol metabolism-related genes are highlighted with rectangles. Expression levels (mean UMI per cell after normalization) for each of the genes in the set are shown at right. **B.** Correlation communities for the same gene set, panel identified using uncorrected PCCs. Note that the target genes are scattered among 5 disconnected communities and are connected with genes with no obvious relationship to cholesterol metabolism. **C**. Positive correlations involving any of the genes belonging to the Reactome Cholesterol Metabolism gene set were divided into “within-community” links and “out-of-community” links. With BigSur (filled symbols), the ratio of within-community to out-of-community links is much higher than with uncorrected PCCs (open symbols), suggesting the latter produce much less enrichment for functionally relevant connections. **D.** Analyses similar to that in panel C were performed for eight additional MSigDB “Hallmark” gene sets. Plotted are the proportions of correlations that are within-group over a range of PCC thresholds. Numbers of genes in each panel are shown in parentheses. **E.** Analyses similar to that in panel D were performed with 150-gene sets chosen at random after first removing genes with mean expression below 0.052 (so that median expression for random genes was similar to that of the functional panel genes.

All detected members of the gene set formed a single network, with many internal positive linkages. Closely connected to this network were eight additional genes (outlined in red) that are not part of the Reactome Cholesterol Biosynthesis set, but belong either to the “Hallmark” cholesterol homeostasis set from MSigDB (*ACSS2, ACTG1, LDLR, SCD, STARD4, TMEM97*); are designated as core cholesterol metabolism genes in recent literature (*ACLY* [68]; *C14or1/ERG28* [69]); or encode sterol binding proteins that serve as feedback controllers of cholesterol biosynthesis (*INSIG1* [70]). Highlighted in orange are two additional genes that regulate lipid biosynthesis more generally, *LPIN1* and *PRRX1*.

Four additional genes are highlighted in gray, as it is possible they are also related to cholesterol metabolism: AKR1A1, CALR, PRDX1, and PMEL. *AKR1A1* encodes an enzyme of the aldo/keto reductase superfamily, which reduces a wide variety of carbonyl-containing compounds to their corresponding alcohols; its orthologues *AKR1C1* and *AKR1D1* are well known to play a role in the metabolism of steroid hormones but *AKR1A1* is thought to act on non-steroid compounds (perhaps this assumption should be revisited). *CALR*, encodes calreticulin, a protein the promotes folding, oligomeric assembly and quality control in the endoplasmic reticulum. As cholesterol synthesis takes place in the ER membrane, it is perhaps not surprising that expression of *CALR* and cholesterol synthesis genes would be co-regulated. *PRDX1* encodes peroxiredoxin, an antioxidant enzyme that reduces hydrogen peroxide and alkyl hydroperoxides and plays a key role in maintaining redox balance. In macrophages, PRDX1 has been shown to be critical for cholesterol efflux during autophagy [71]. *PMEL* encodes a major component of the melanosome. Interestingly, it has been reported that cholesterol strongly stimulates melanogenesis in melanocytes and melanoma cells [72].

In addition to these genes, a set of tightly interconnected genes that have no known association with cholesterol metabolism is peripherally connected to this community (outlined by a dotted line in Figure 6A). A large proportion of these genes encode proteins associated with the cell cycle, especially with functions associated with mitosis, such as mitotic checkpoint events, and cyto- and karyokinesis. Strongly connected to these mitotic genes is *LBR*, which encodes the Lamin B receptor. LBR is a multifunctional protein: it plays a key role in cholesterol biosynthesis, catalyzing the reduction of the C14-unsaturated bond of lanosterol, but also anchors the nuclear lamina and heterochromatin to the inner nuclear membrane, and in so doing is strongly regulated by phosphorylation during the cell cycle [73]. A functional link between expression of cholesterol biosynthesis genes and mitosis may arise from that fact that cholesterol synthesis normally rises dramatically during G1 phase and if prevented from doing so will result in G1 arrest [74]. It thus may make sense to upregulate the production of cholesterol biosynthetic genes as cells go into mitosis, so they are available to act in the G1 phase that immediately follows. To the right of the graph in Figure 6A the genes of the Reactome Cholesterol Biosynthesis set are listed alongside their relative expression levels (UMI/cell after normalization) in this data set. Highlighted in yellow are those for which BigSur identified significant positive correlations, nearly all of which were toward the higher end of expression levels. These results illustrate an important limitation of BigSur, which is that statistical power depends on the number of non-zero entries in the data, which will, in turn, reflect expression level, sequencing depth, and the number of cells analyzed. Examination of the raw data indicates that the point at which BigSur begins to fail to identify significant correlations occurred, for these genes, when the total number of non-zero entries (among 1582 cells) fell to about 320; this corresponded to a mean expression level of about 0.3 UMI/cell. For more strongly correlated genes one should of course expect fewer entries to be necessary to achieve significance, but based on the results here we suggest that, when seeking to identify the kinds of weak correlations associated with noise-coupled gene-regulatory networks, a minimum of several hundred non-zero entries per gene may be a reasonable threshold. In practice, one should be able to adjust the number of genes that fulfill this criterion either by varying the number of cells analyzed or varying the depth of sequencing (or both).

Fig. 6B plots the results of the identical exercise as in Fig. 6A, carried out using uncorrected PCCs obtained directly from normalized gene expression data, rather than using BigSur. A threshold of PCC = 0.29 was selected so that the number of Reactome Cholesterol Biosynthesis genes that exhibited above-threshold correlations was the same as in Fig. 6A. In this case, however, the cholesterol biosynthesis genes did not form a single community, but rather separated into 5 disconnected groups. Most groups contained links to many genes, none of which appeared, on inspection, to bear any relationship to cholesterol metabolism. These data support the view that most of the links detected using uncorrected PCCs are false positives.

One way to extend this analysis to many other functional gene sets is suggested by the observation that, among the correlations in Fig. 6A involving Reactome Cholesterol Biosynthesis genes, 25 occur between member genes (“within-community”) while 157 involve other genes (“out-of-community”). In contrast, in Fig. 6B there are 143 out-of-community links and zero in-community ones. Fig. 6C shows how the numbers of within-community and out-of-community links change as the PCC significance threshold is changed, both for Big Sur and uncorrected PCCs. These results suggest that the fraction of all correlations that is in-community can be used as a measure of the specificity with which functionally relevant correlations are identified by any given method. In Fig. 6D, we used this approach to contrast the performance of BigSur (filled symbols) with uncorrected PCCs (open symbols) for 8 different gene sets, each of which contained between 49 and 200 members. Using BigSur, the proportion of links that were in-community varied between ∼1% and 30% depending on the gene set, usually increasing as the equivalent PCC threshold was made more stringent. Using uncorrected PCCs, the proportion of in-community links varied between 0.2% and 1% and did not change appreciably with PCC threshold. For comparison, Fig. 6E performs a similar analysis using 8 “random” gene sets—sets of 150 genes randomly selected from genes with similar overall expression to those in the gene sets used in Fig. 6D. The results suggest that the performance of BigSur on functionally related genes is far above chance level, while uncorrected PCCs perform at or around that level.

## Discussion

The above results suggest that *bona fide* communities of co-regulated genes can be identified with high specificity by carefully mining weak correlations within groups of a few thousands of relatively homogeneous cells. BigSur achieves the accuracy to do this first by correcting measures of correlation for unequal sequencing depth and the added variance contributed by gene expression noise, and subsequently by estimating an individual *p*-value for each gene pair—thereby overcoming the strong effect of gene expression distribution on the likelihood of a correlation coefficient arising by chance. In contrast, the use of uncorrected PCCs to identify co-regulated genes performed poorly on both synthetic and real scRNAseq data.

In developing BigSur, we sought to avoid normalization steps and the use of expression thresholds and minimize user-provided parameters to the greatest extent possible. The major user input to BigSur, besides a UMI matrix, is a coefficient of variation for gene expression noise, *c*, which can be quickly estimated by fitting a plot of the modified corrected Fano factor against gene expression level (Fig. 2B). In reality, the magnitude of the noise of gene expression may be different for different genes, and for some the Poisson log-normal distribution may not be the best approximation of the noise. Although these factors likely degrade the performance of BigSur, we note that the value of *c* only significantly impacts the highly expressed genes, for which (due to low sparsity) the detection of significant correlations is a less challenging task.

A more subtle source of potential error comes from fact that, In calculating modified corrected Pearson residuals, the value of 𝜇_*ij*_ used by BigSur is determined empirically from a finite set of cells, i.e., it is an estimator of 𝜇_*ij*_. Furthermore, whereas it is an unbiased estimator when *µ* is Poisson-distributed, this is not generally the case for more skewed distributions, such as Poisson-lognormal [43]. It is unknown whether these sources of error have much impact on the determination of correlation coefficient *p*-values by BigSur, and additional work will be necessary to investigate this question.

Despite these concerns, the ability of BigSur to identify gene communities that are closely related in function (Table 1), as well as add new, functionally related genes into known gene sets (Fig. 6), suggests that it already operates at a level of performance that can be useful to cell and tissue biologists. Particularly interesting are the questions it raises about linkages within communities—for example, why do genes encoding cystatin endopeptidase inhibitors (*CST1*, *CST4*, *CSTB*) correlate with genes involved in glycolysis and cellular respiration (community E)? Why do genes encoding protein chaperones correlate with genes involved in RNA processing (community G)? Why do genes involved in Golgi and lysosome function correlate with genes related to hypoxia (community J)?

The data in Table 1 also suggest that new cell biology may be discovered by examining genes labeled “not accounted for”, i.e., genes that are associated with a community but do not, as a group, overlap substantially with any mSigDB dataset. For example, 38 of the 213 genes that correlate with cell cycle community B are not currently annotated as cell-cycle related. Among the strongly-coupled unfolded-protein response genes in community C, one also finds genes encoding secreted (*GDF15, PSAP, SPARC*) and cell surface molecules (*DDR2*, *GPNMB*, *ITGA4, IL1RAP, PMP22, SLC1A5*); the potential relationship of these genes to cellular stress responses may deserve heightened attention.

Indeed, each of the communities in Table 1 suggests new and unexpected forms of co-regulation of gene expression. It will be interesting to see how many of these are reproduced in other cell types—a task that can be efficiently approached by applying BigSur to the large number of already existing scRNAseq data sets.

It is instructive to note that few of the gene-gene relationships detected by BigSur could have been revealed by the typical analytical steps of cell clustering and identification of differentially expressed genes, as most of the gene communities identified here are not associated with sufficient total variation in gene expression to be useful drivers of cell clustering. On the other hand, clustering did play an important role here in reducing the heterogeneity of the sample to which BigSur was applied. Although most of the gene communities that were identified using the 1,582-cell subcluster 1.2 were also observed when BigSur was applied to the entire sample of 8,640 cells (not shown), clusters were more difficult to visualize and analyze in the latter case, thanks to a large background of cell-type (or cell-state)-specific gene expression (which generated numerous additional correlations). This experience suggests that iterative application of BigSur analysis and clustering (potentially using correlated gene communities as features) can provide a useful pipeline for identifying meaningful gene communities of manageable size.

It is interesting that one of the strongest axes of variation we detected in this study involved mitochondrially-encoded genes and genes coding for ribosomal subunits, with both communities strongly anti-correlating with each other (both before and after “damaged” cell removal and subclustering). Because the mitochondrial community consists only of mitochondrially-encoded, and not cytoplasmically-encoded, mitochondrial genes, the simplest interpretation is that this community reflects cell-to-cell variation in the number of mitochondria (or, more accurately, mitochondrial genomes). What then might explain anti-correlation between mitochondrial number and transcripts for ribosomal proteins? Given that protein synthesis requires both specialized machinery (ribosomes) and a source of energy (mitochondria), one might expect to see positive, rather than negative, correlation between the agents that orchestrate these processes. Yet this intuition is correct only in a long-time-averaged sense, and does not necessarily apply if demand for protein synthesis fluctuates. In mammalian cells, ribosomal protein mRNAs are long-lived, with half-lives in the 5-10 hour range [75], whereas mitochondria can replicate on a time scale of 1-2 hours [76]; accordingly, one process may systematically lag the other, producing the kind of anti-correlation we observe here. While this interpretation is speculative, it demonstrates how the analysis of gene-gene correlations can motivate new hypotheses about transcriptome-scale regulation of cell biology.

Under the expectation that most networks of gene correlation reflect shared transcriptional regulation, we might have expected to identify upstream transcription factors more frequently in most of the networks we discovered. In some cases, this clearly did occur: For example, the transcription factors *ATF4*, *AT6*, and *XBP1*, which were detected in community C, are known controllers of the unfolded protein response, the components of which show up in the same community. Transcription factors related to cell cycle progression and DNA damage-repair strongly associated with cell-cycle communities A and B. *ZKSCAN1*, a transcription factor targeted to mitochondria [77] associates with the mitochondrially-encoded gene community H. However, in many cases, expected transcription factors (e.g., *SREBF1* or *SREBF2* in community K) were not observed. This may reflect limitations in statistical power, as transcription factor genes tend to be expressed at a somewhat lower level than other genes—although in this data set expression of the average observed transcription factor was only about half that of the average observed gene. Another likely explanation is that gene regulation is often achieved through the post-translational modification of transcription factors, rather than regulation of their mRNAs. Alternatively, it may reflect the importance of factors that act post-transcriptionally, such as miRNAs (which are not detected here), in controlling gene expression. In future, it will be important to extend BigSur to take account not only of miRNAs but also of “multi-omic” features, such as chromatin accessibility.

Although we have focused here primarily on the use of BigSur as a tool for discovery of gene-gene correlations, it is worth pointing out that the intermediate steps in the BigSur pipeline produce useful tools for other analytical procedures, some of which were exploited here. For example, methods for feature selection for cell clustering commonly involve thresholds (e.g., expression levels) and cutoffs (e.g., numbers of features) that are arbitrary, and may not be ideal choices for every data set. Use of the modified corrected Fano factor 𝜙′, and its associated *p*-values, can provide a less arbitrary approach to feature selection, which can outperform other methods on challenging tasks, such as finding rare cell types, or subclustering cells that differ only modestly [78]. In addition, as we did here when dividing cells into subclusters based on the expression of ribosomal and mitochondrial genes (Fig. 4), one may also use communities of correlated genes themselves as features for clustering—in this way leveraging not just variation but co-variation to drive clustering. Finally, whereas it is common practice to cluster cells based on their normalized expression values, the matrix of modified corrected Pearson residuals that BigSur calculates almost certainly provides a more accurate starting point for clustering, as it avoids artifacts introduced by normalization, and corrects for the inflated variance associated with highly expressed genes.

## Conclusions

It has long been suspected that, by mining gene expression correlations within single cell transcriptomes, it should be possible not only to identify groups of genes that distinguish among cell types, but also discover small-scale gene regulatory networks that operate within cell types. We have provided evidence here that this goal can be achieved, first by modifying and correcting the metric of correlation and then using an analytical approach to ascertain statistical significance for every gene pair. When applied to scRNAseq data, this dramatically reduced the number of false positives that would have been identified using traditional methods, and enabled the identification of biologically relevant gene communities, both known and novel.

## Methods

As the first step in calculating 𝜙, and PCC′, we begin with “raw” (neither normalized nor log-transformed) UMI data and calculate, for each gene in each cell, a cell- and gene-specific Pearson Residual, defined according to eq. 1. To do so, we first calculate a cell- and gene-specific expected value (𝜇_*ij*_) which we obtain for each gene by averaging its values over all cells, then scaling that in each cell by to the relative proportion of total UMI that each cell contains. This is essentially the same procedure used in the simplest form of normalization, except that, rather than normalize the data matrix, we are normalizing the term for the mean in the Pearson residual.

Next, each modified Pearson residual is divided by 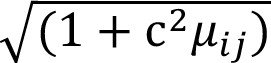, where *c* is a constant between 0.2 and 0.6. Ideally, *c* should be chosen individually for each gene, depending on prior knowledge of the level of gene expression stochasticity, however, in the absence of prior knowledge we typically find *c* empirically (see Fig. 2B). It should be noted that, for many scRNAseq data sets, most values of 𝜇_*ij*_ will be < 1, meaning the effect of the choice of *c* on most of the data is often relatively small. To obtain 𝜙, for each gene, the Pearson residuals for each cell are squared, summed, and divided by *n*-1, where *n* is the number of cells.

To calculate *p*-values for 𝜙, any given gene, we consider the null hypothesis to be the expected number of transcripts in each cell is 𝜇_*ij*_, i.e., the total number of transcripts across all the cells partitioned in proportion to the number of total UMI in each cell. As noted above, because BigSur determines the value of 𝜇_*ij*_ empirically—by summing up genes UMI across all cells and multiplying by the fraction of total UMI in each cell—𝜇_*ij*_is technically an estimator of the cell-specific expectation value for each gene and cell.

To calculate *p* we need to know how the sums of squared Pearson residuals should be distributed for any given set of 𝜇_*ij*_ and *c*. As discussed above, we take 𝜇_*ij*_to have a Poisson-log-normal distribution, allowing us to calculate the moments of the distribution of squared Pearson residuals from the moments of the Poisson and log-normal distributions, and from there the moments of the distribution of sums of squared Pearson residuals. In the end we obtain, for each gene *j*, a finite set of moments (typically 4 or 5) for the distribution of 𝜙,that would be expected under the null hypothesis for that particular gene, given the values 𝜇_1*j*_, 𝜇_2*j*_, 𝜇_3*j*_ … 𝜇_*nj*_ and *c*. We then use Cornish-Fisher approximation of the Edgeworth expansion [79] as a reasonably computationally efficient way to approximate the *p*-value associated with any given observation of 𝜙,, given that distribution.

The procedure for obtaining PCC′ proceeds in the same fashion, starting with the same modified corrected Pearson residuals, but now taking the dot product of the vectors of Pearson residuals for each pair of genes, and dividing by *n*-1 times the geometric mean of the 𝜙,values for those genes (eq. 2). Moments of the expected distributions of PCC′ are calculated analytically in exactly the manner described above, with the slight complication that the 𝜙, terms in the denominator are not strictly independent of the Pearson residuals in the numerator, but to a good first approximation may be treated as such, as they aggregate information across all the Pearson residuals. The Cornish-Fisher approximation is again used, as described above, to assign *p*-values to both tails of the resulting distribution. In solving the 4^th^ degree polynomial equations produced by this method (a computationally slow step), we improve speed by sacrificing accuracy specifically for those *p*-values that clearly fall below thresholds of interest.

Because the number of possible gene-gene correlations scales roughly as the square of the number of genes, the need to correct for multiple hypothesis testing is particularly acute. Given that the observations are not independent from one another, Bonferroni correction is clearly too conservative, and thus we use the Benjamini-Hochberg algorithm [49] for controlling the false discovery rate.

### Simulating scRNAseq data

In Figure 2 we simulate the expression of 1000 genes across 999 equivalent cells, where by equivalent we mean that, for each gene, a single “target” transcript level was chosen from a log-normal distribution, the mean of which was selected so that the logarithm of its value varied evenly across the gene set, over the range from 0.035 to 3467 transcripts per cell. The coefficient of variation of each log-normal distribution was taken to be 0.5 for all genes. Next, we generated a set of 999 scaling factors, drawn from a log-normal distribution with mean of 1 and coefficient of variation of 0.75, selected so that the logarithm of the value varied vary smoothly across the range. Finally, we generated a UMI value to each gene in each cell that was a random variate from a Poisson distribution with a mean equal to the target for that gene times the scaling factor for that cell. The result was a set of gene expression vectors of length 999, with mean values varying between 0.001 and 231.

### Analysis of melanoma cell line data

Data from droplet-based sequencing of subcloned WM989 melanoma cells (GEO accession GSE99330), which had been pre-processed to remove UMI judged not to be associated with true cells, were imported and further pre-processed in the following way: First we removed all known pseudogenes (comprehensive lists of human pseudogenes were obtained from HGNC and BioMART). Pseudogenes derived by gene duplication or retrotransposition are often highly homologous to their parent genes, creating ambiguity in the mapping of the short-read sequences used in scRNAseq. When pseudogenes were not removed from analysis, we frequently detected strong correlations between pseudogenes and parent genes that very likely represented the effects of ambiguous mapping, rather than true correlation. After pseudogene removal, the number of detected genes was 27,526. As both theory (Fig. 1) and experience indicated that statistically significant correlations were usually unobservable for genes with UMI in fewer than 0.001% of cells, we further eliminated genes expressed in fewer than 8 of the 8640 cells; this reduced the size of the gene set to 17,451, and the number of possible unique correlations to 152,259,975 (while not strictly necessary, this step reduces computational time by ∼2.5 fold, since the number of correlations to test varies approximately quadratically with the number of genes).

### Calculating paralog pair and protein-interaction enrichment scores

A curated list of 3,132 paralogous pairs of human genes was downloaded from [80]. A list of physical human protein-protein interactions was downloaded from BioGrid (https://downloads.thebiogrid.org/File/BioGRID/Release-Archive/BIOGRID-4.4.218/BIOGRID-MV-Physical-4.4.218.tab3.zip) and supplemented with additional data from HIPPIE v2.3 (http://cbdm-01.zdv.uni-mainz.de/~mschaefer/hippie/), to produce a list of 790,008 gene pairs. To calculate enrichment, we first calculated the fraction of statistically significantly positively correlated gene pairs identified by BigSur that overlapped with either the paralog-pair or protein-protein interaction pair database. Next, we removed from the databases all gene pairs involving genes not detected in the scRNAseq data and divided the remaining number by the total number of possible gene-gene correlations (i.e. *m*(*m*-1)/2, where *m* is the number of genes in the scRNAseq data) to yield the expected frequency of paralogous or interacting pairs under the hypothesis they are randomly distributed among all possible pairwise correlations. The ratio between the observed frequency and expected frequency was considered to be the fold enrichment.

### Extracting (and pooling) gene communities

BigSur generates a matrix in which rows and columns are genes, and entries are signed equivalent PCCs—which are derived by using the inverse of the Fisher formula on the *p*-values returned by BigSur, together with the signs of the values of PCC′. Although it is a derived quantity, the equivalent PCC is a useful form in which to store correlation data, not only because it is signed, but also because it adjusts for differences in data length (number of cells), so that similar “strengths” of correlation would be expected to translate into similar equivalent PCCs, even across samples with very different numbers of cells (cell number has a large effect on the relationship between correlation strength and *p*-value).

Only equivalent PCCs that were judged statistically significant according to a user-supplied threshold (e.g., a Benjamini-Hochberg FDR) were included, all others being set to zero. This matrix was converted to an unweighted adjacency matrix (all nonzero values replaced with 1) and the *walktrap* algorithm (with a default setting of 4 steps) was used to identify initial communities [52]. Because this produces communities connected by both positive and negative links, each community was then subjected to a second round of community-finding, after first setting negative links to 0, thus allowing subcommunities that negatively correlate with each other to be separated.

To identify instances in which walktrap had subdivided communities too finely, we manually examined the number of positive correlations between genes in each community and each other community, recursively merging communities in which the number of inter-community correlations was particularly large (compared with the number of possible links between communities). In addition, in rare cases in which communities returned were very large (e.g., in the thousands), we subdivided them by applying walktrap an additional time.

### Cell clustering based on correlated features

Feature selection refers to the process of identifying genes that capture important dimensions of variation on which cells may be clustered. A variety of approaches have been proposed for identifying such genes, and many work equivalently under most circumstances (with tens of thousands of genes, clustering is often a highly over-determined problem). Under challenging circumstances (e.g. when the number of true difference separating clusters is small, or the number of cells in a state is small), we have shown [78] that 𝜙′ is a measure of variability at least as good as others, and because Big Sur returns both 𝜙′ and *p*-values, one may avoid selecting too large a set of features (which can defeat clustering algorithms).

It has also been pointed out, however, that not only the statistical features of individual genes, but also their interdependencies (i.e., correlations) should ideally be used to inform clustering [24]. We recognized that the communities identified by BigSur represent ideal sources of features, particularly if we emphasize those community members that are the most highly connected to each other. We also recognized that the modified corrected Pearson residuals generated by BigSur provide a more sensible set of vectors to use as the input to clustering than either raw or normalized UMIs (for the same reasons that 𝜙′ and PCC ′ are improvements over their unmodified, uncorrected forms). Using this approach, we routinely found that well defined clusters can often be reliably obtained using sets of highly connected genes as small as 50-75 each. This approach was used to repetitively subcluster the scRNAseq data from WM989 melanoma cells, at each step running Big Sur and using the 50 most connected genes of the clusters containing the majority of ribosomal genes, and the 50 most connected genes of those containing the majority of mitochondrially encoded genes, as features.

### Graphical display of correlations

Matrices representing statistically significant correlations were plotted using the *GraphPlot* function of Mathematica software, in which vertices were arrayed either by Spring Embedding or Spring Electrical Embedding. Edges were colored green when positive and red when negative. Vertex locations were first determined according to the graph produced after deleting negative edges, after which vertices connected only by negative edges were added in. Symbols used to represent vertices were scaled so that their areas were proportional to the mean expression level of the gene represented. Correlation strengths (the absolute values of the equivalent PCCs) are not represented on these images.

## Code Availability

R and Mathematica implementations are available under the Lander lab profile on GitHub (https://github.com/landerlabcode/).

## Abbreviations

BigSur, Basic Informatics and Gene Statistics from Unnormalized Reads; CV, Coefficient of variation; ER, Endoplasmic reticulum; FDR, False discovery rate; PCA, Principal component analysis; PCC, Pearson correlation coefficient; PCC’, Modified, corrected Pearson correlation Coefficient; ROC, Receiver operating characteristic; scRNAseq, Single cell RNA sequencing; UMAP, Uniform manifold approximation and projection; UMI, Unique molecular identifier.

## Declarations

Ethics approval and consent to participate: Not applicable. Consent for publication: Not applicable.

## Availability of data and materials

The datasets generated and code used for analysis during the current study are available in the BigSurM Github repository, https://github.com/landerlabcode/BigSurM/tree/main/Datasets%20and%20Analyses

## Author contributions

KS conceived of the work, developed and implemented code, was responsible for posting code and data to github, and contributed to writing of the manuscript. ED conceived of the work and contributed to writing of the manuscript. JG helped conceive of the study and contributed to the development of code. SA provided financial support and edited the manuscript. QN provided financial support, advice on code development and edited the manuscript. AL conceived of the work, developed and implemented code, and contributed to writing of the manuscript. All authors read and approved the final manuscript.

## Supporting information

Mathematical Appendix

## Acknowledgements

We thank Weining Shen (UCI) for advice on statistical analysis. We acknowledge Mika Caldwell and Yilun Zhu for helpful feedback and testing of code.

## Figure legends

**Table 2.** Mitochondrially-encoded and ribosomal protein gene communities identified following unsupervised clustering of each of the four cell clusters, 1.1, 1.2, 2 and 3. Mitochondrially-encoded and ribosomal protein genes are highlighted in bold.

**Figure S1. Significance of uncorrected Pearson correlation coefficients (PCC), as calculated by BigSur versus the Fisher formula, binned by gene expression.** scRNAseq data were as described in Figure 3. Data points representing pairs of genes were divided into 21 bins based on the mean expression levels of each gene, and the results for each bin were plotted as described in Fig. 3C. The abscissa shows PCC (calculated from default-normalized data) while the ordinate gives the negative *log_10_* of *p*-values determined by BigSur, i.e., larger values mean greater statistical significance. Orange and gray shading indicate gene pairs judged significant by BigSur (FDR<0.02). Blue and orange show gene pairs that would have been judged statistically significant by applying the Fisher formula to the PCC, using the same *p*-value threshold as used by BigSur. The blue region contains gene pairs judged significant by the Fisher formula only, while the unshaded region shows gene pairs not significant by either method. Numbers in the lower right corner of each panel are the total numbers of possible correlations (blue), statistically significant correlations according to the Fisher formula (green), and statistically significant correlations according to BigSur (red).

**Figure S2. Significance of modified corrected Pearson correlation coefficients, (PCC ′), as calculated by BigSur versus the Fisher formula, binned by gene expression.** scRNAseq data were as described in Figure 3. Data points representing pairs of genes were divided into 21 bins based on the mean expression levels of each gene, and the results for each bin were plotted as in Fig. S1. The inset compares the PCC′-*p*-value relationship determined by BigSur (for genes with relatively high expression levels) with that predicted by the Fisher formula (dashed red line), showing that, for highly expressed genes, the two methods agree well.

**Figure S3. Distribution of equivalent PCCs.** The *p*-values obtained by BigSur for the melanoma cell line were transformed using the inverse of the Fisher formula to a set of “equivalent” PCCs. Since the Fisher formula operates on the absolute values of correlations, each calculated equivalent PCC was assigned the sign of the PCC′ value for the same gene pair. Equivalent PCCs provide a measure of correlation strength that can be compared across data sets with differing numbers of cells. They may be understood as a measure of how strongly correlated two normally distributed vectors (of any given length) would need to be to produce the observed p-value. Here, only those gene pairs judged significant by BigSur are shown. The fact that so many weakly correlated gene pairs (|equivalent PCC| <0.05) are nevertheless statistically significant is a function of the long vector length in this experiment (>8000 cells).

**Figure S4.** Gene communities A and B (see Table 1) from cell cluster 1.2. Green edges depict significant correlations. Transcription factor vertices are displayed as yellow boxes with gene names in blue. In the boxed insets the same graphs are overlayed in brown to highlight links supported by known protein-protein interactions.

**Figure S5.** Gene communities C, and D (see Table 1) from cell cluster 1.2. Genes and links are highlighted as in Fig. S4

**Figure S6.** Gene communities E, F, G and H (see Table 1) from cell cluster 1.2. Genes and links are highlighted as in Fig. S4.

**Figure S7.** Gene communities I, K, L and M (see Table 1) from cell cluster 1.2. Genes and links are highlighted as in Fig. S4. In communities L and M, links supported by known protein-protein interactions are highlighted in brown.

## Additional Files

Additional File 1: Table 1. Gene communities identified in cell subcluster 1.2

Additional File 2: Table 2. Mitochondrially-encoded and Ribosomal Protein gene communities identified in each of the four cell clusters.

Additional File 3: Appendix (Mathematical derivations underlying the calculations of statistical significance by BigSur).

Additional File 4: Supplemental Figure S1

Additional File 5: Supplemental Figure S2

Additional File 6: Supplemental Figure S3

Additional File 7: Supplemental Figure S4

Additional File 8: Supplemental Figure S5

Additional File 9: Supplemental Figure S6

Additional File 10: Supplemental Figure S7

**Table S1.**
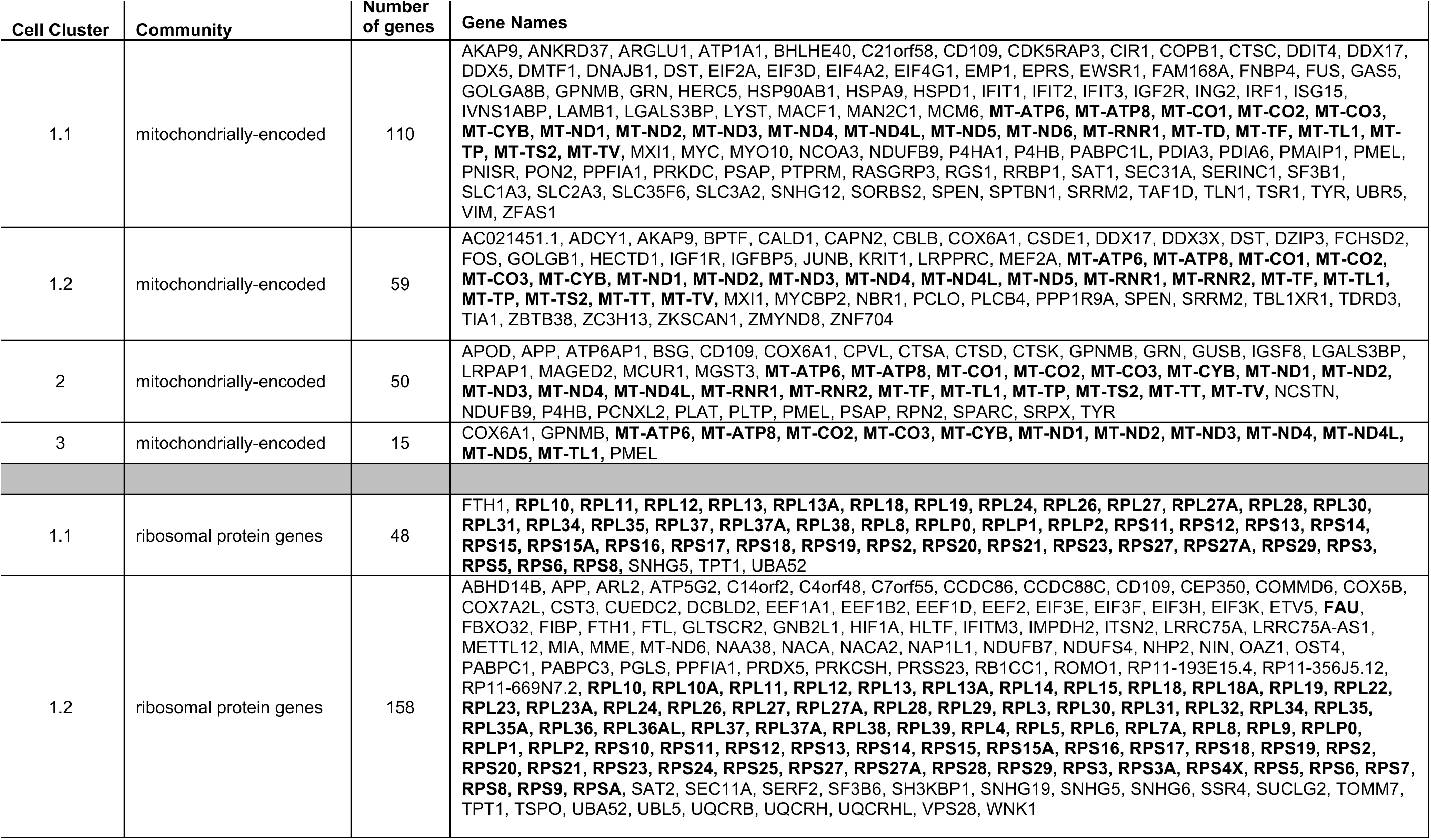
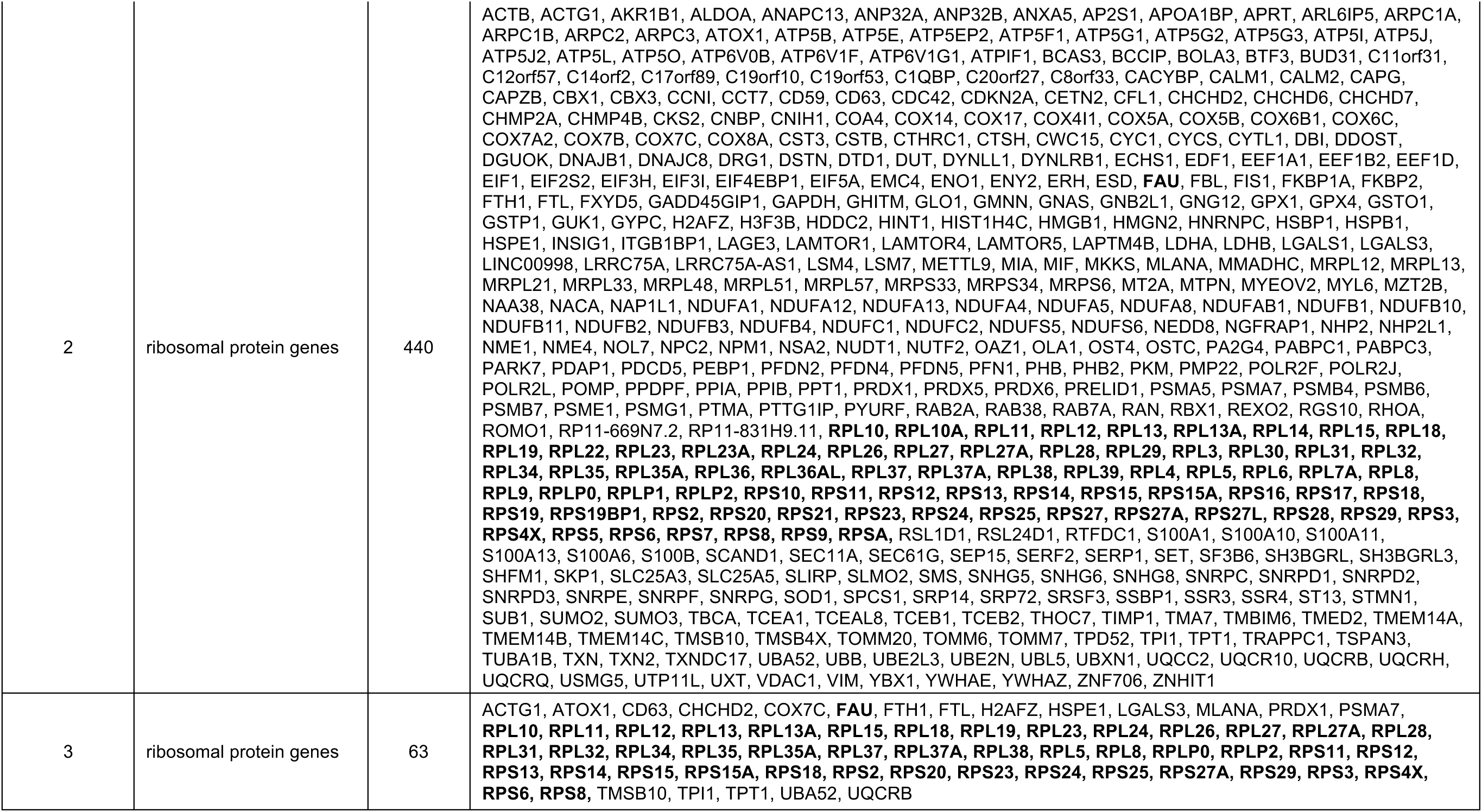
Mitochondrially-encoded and Ribosomal Protein gene communities identified in each of the four cell clusters.

