## Supplementary figures and images for "Leveraging gene correlations in single cell transcriptomic data"

### Supplemental Figures S1-S7

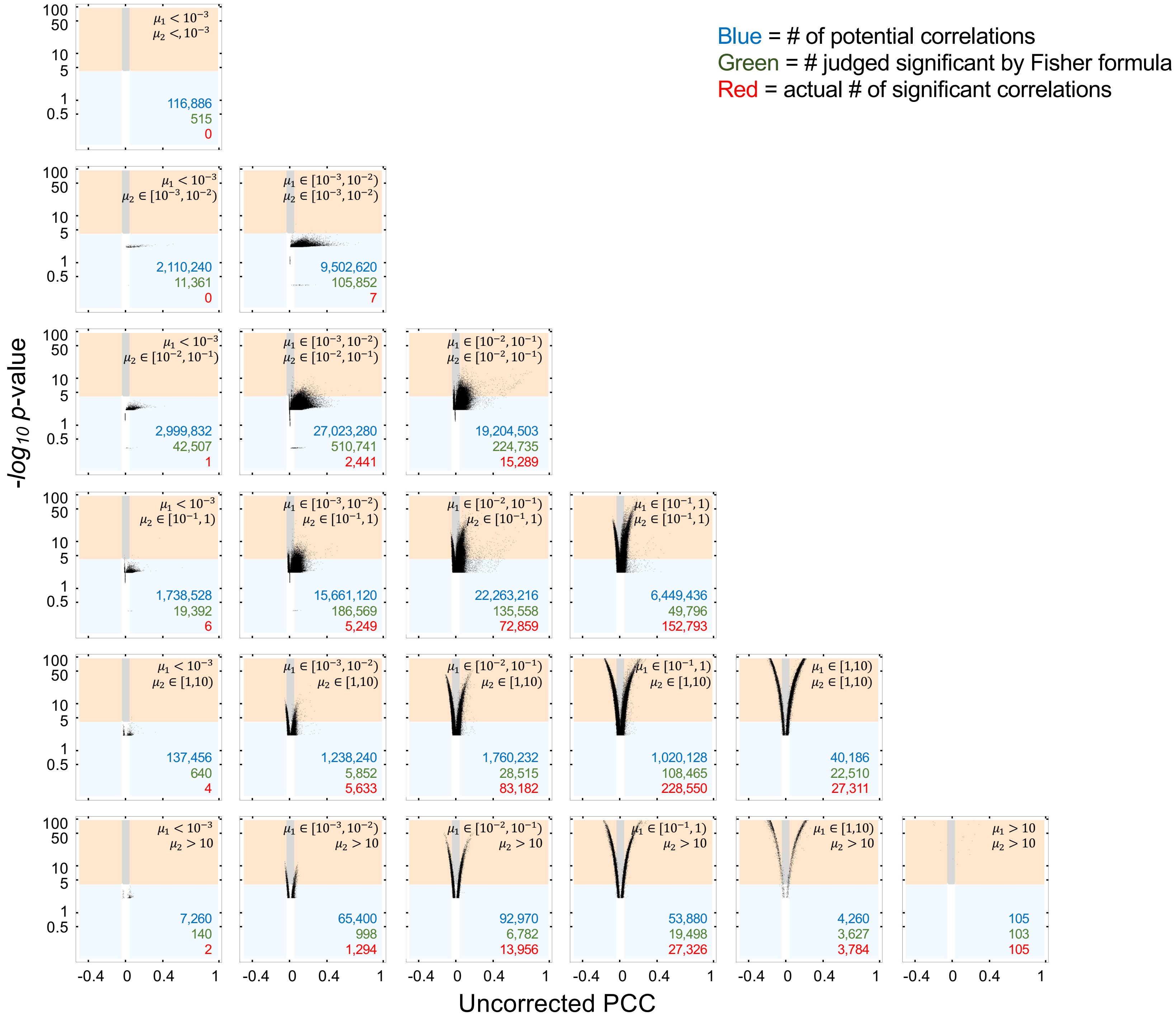

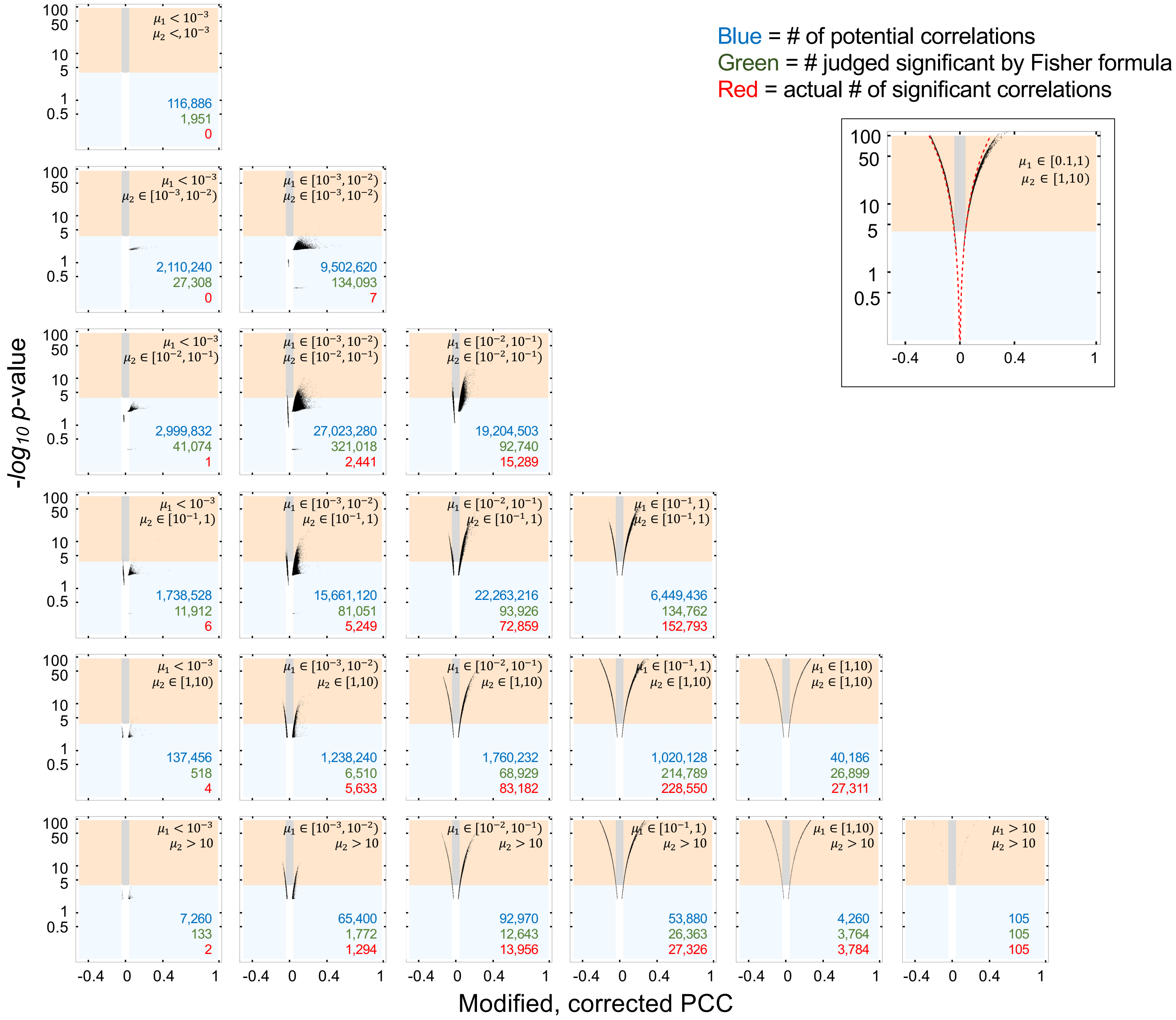

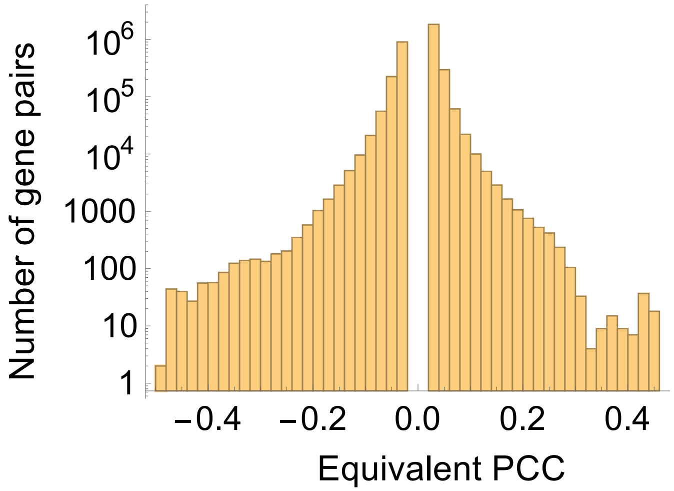

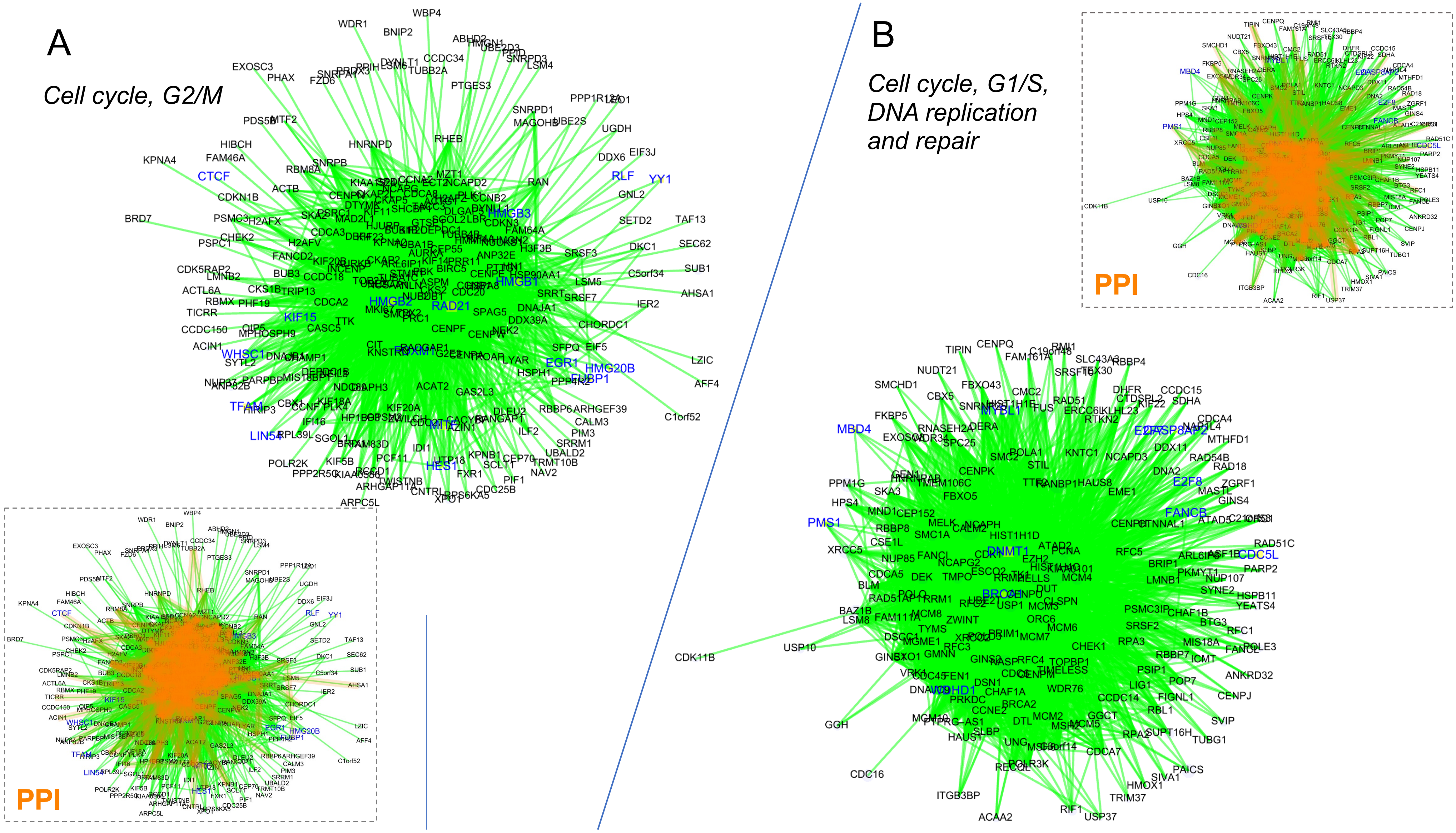

C

*Unfolded protein response, ER stress*

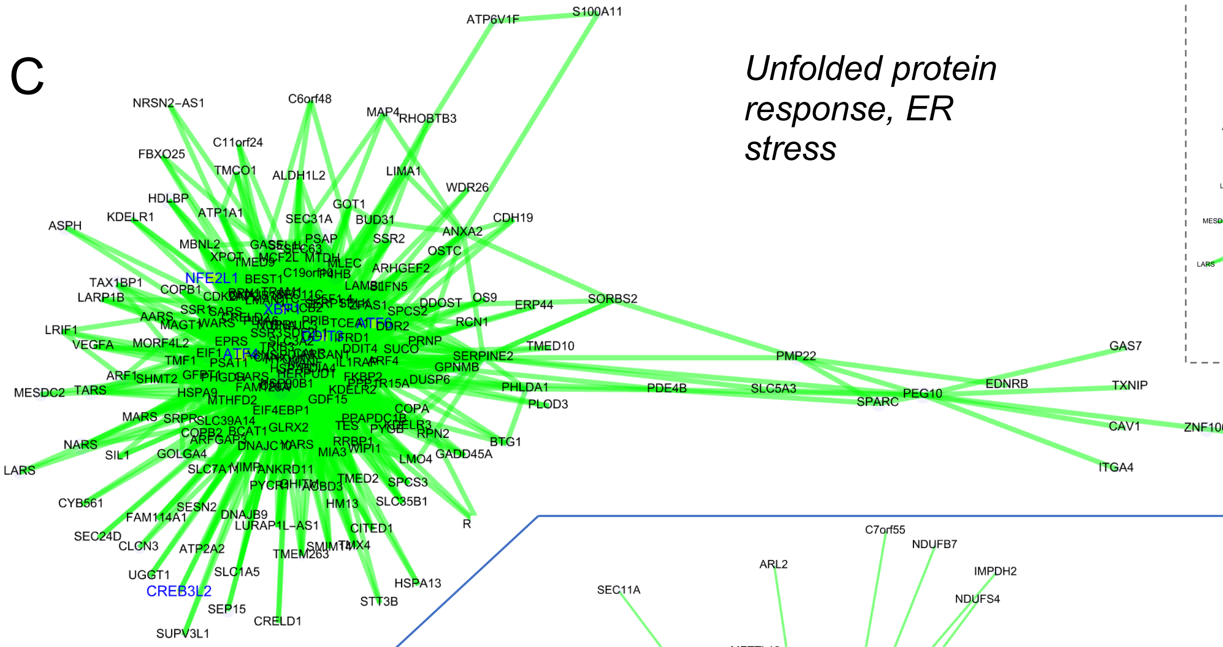


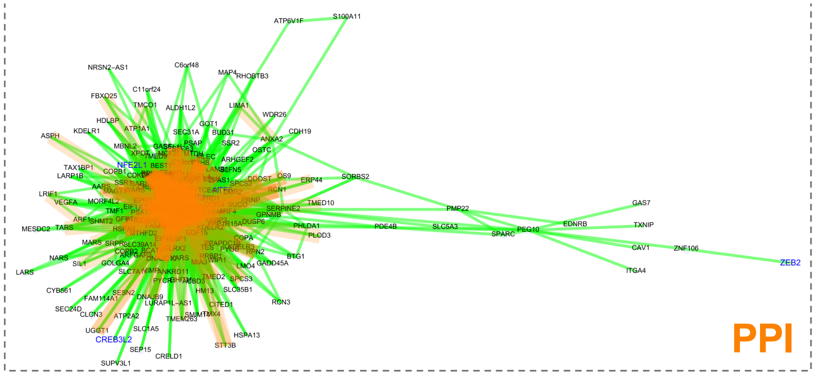

D

*Ribosome biogenesis, protein translation*

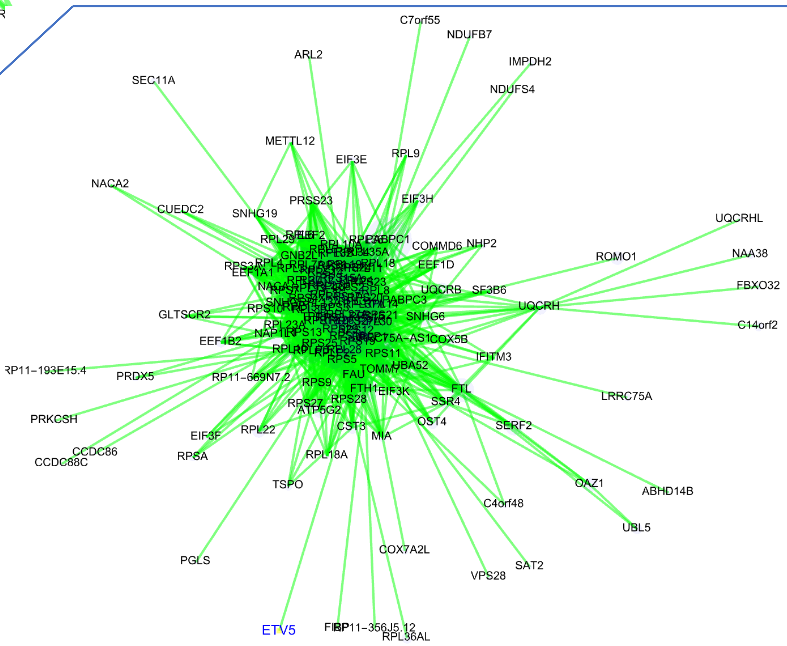


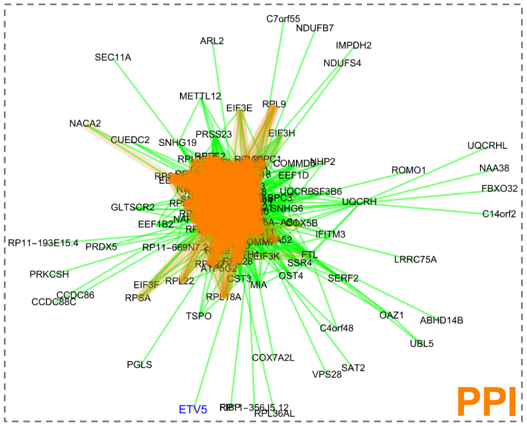

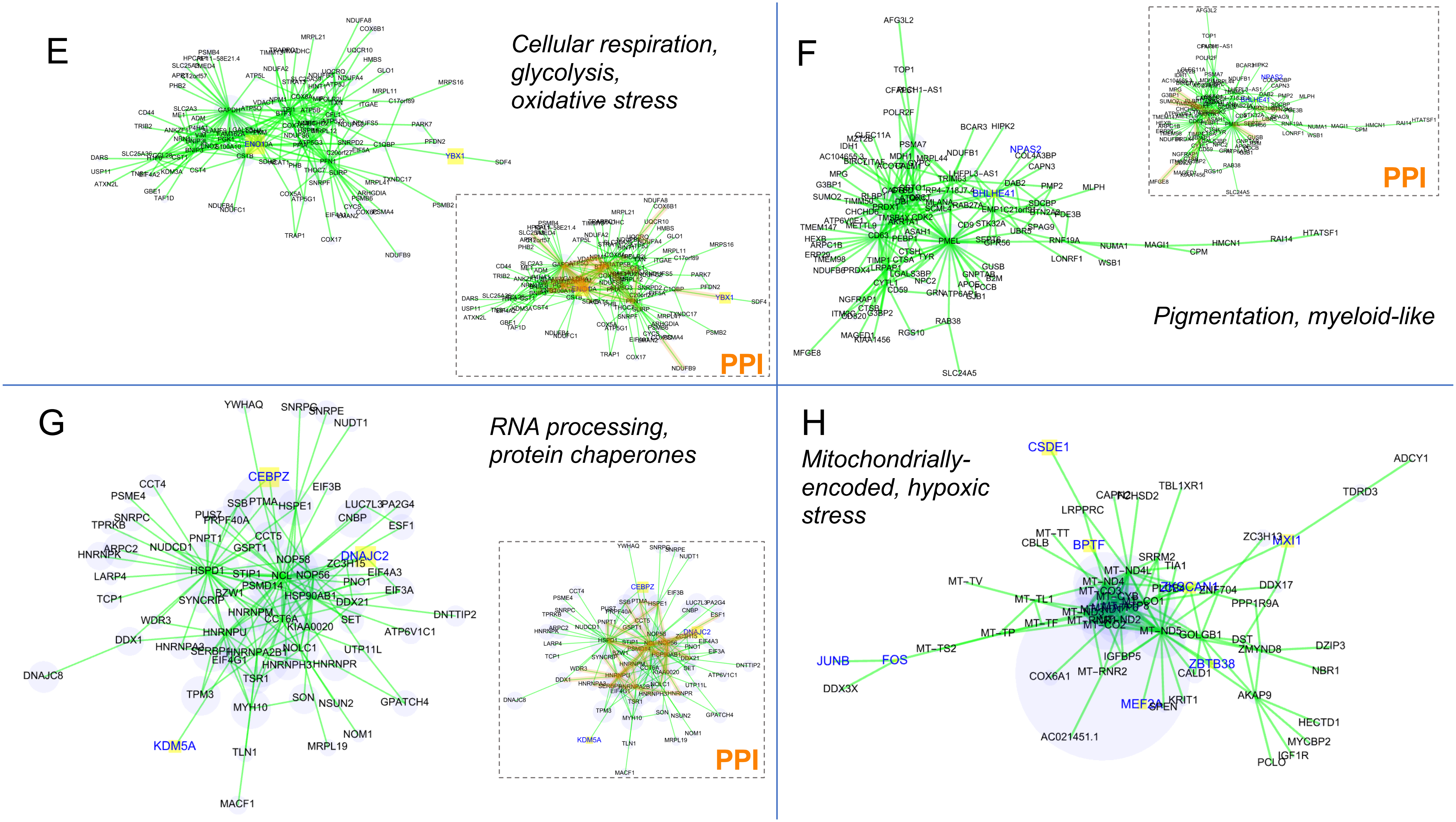

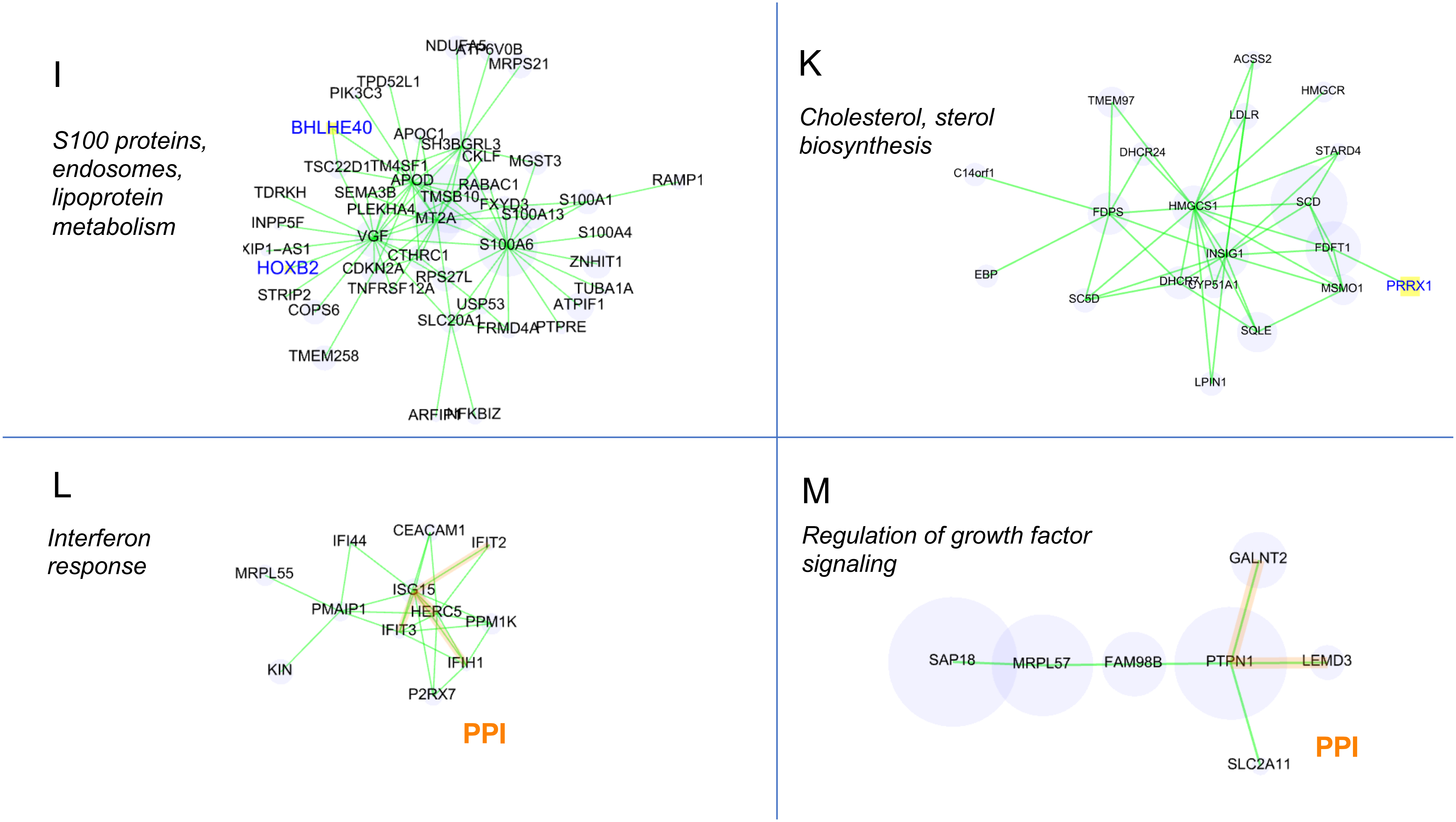
