## Supplementary material for "Leveraging gene correlations in single cell transcriptomic data": Mathematical Appendix

#### Deriving p-values from expectation matrices

Let $a_{ij}$stand for the UMI entries of the *m* x *n* matrix of *m*-genes and *n*-cells. As described above, we use the totals over all i and the totals over all j to generate an "expectation matrix", i.e. a matrix whose entries $\mu_{ij}$correspond to the expected values of the UMI for each gene in each cell under the null hypothesis (the hypothesis that gene expression is identical across cells).

To derive *p*-values for modified corrected Fano factors ($\phi$') or modified corrected Pearson Correlation Coefficients (PCC') we must derive, starting from the values in the expectation matrix and the expected distribution of those values, the expected distributions of the $\phi$' and PCC' under the null hypothesis. Identifying those distributions enables one to determine the probability, under the null hypothesis, that any given observation would have occurred.

The way this is done in two steps: First we calculate the moments (or equivalently, cumulants) of $\phi$' and PCC' under the null hypothesis. Second, we use the Cornish Fisher method to reconstruct the tails of distributions from their cumulants, determining how far out on the tail any empirical observation lies.

#### The Poisson-Log-Normal (PLN) Distribution

We define the Poisson Log Normal distribution as the discrete distribution obtained by random sampling of integer values from a Log normal distribution. Specifically, we parametrize the Log normal distribution in terms of its mean $\mu$ and coefficient of variation c (its actual mean and coefficient of variation, not that of the underlying normal distribution in terms of which log normal distributions are often defined). According to Bulmer (1974), compounding the the Poisson distribution with any other distribution that has a PDF of f($\lambda$) gives a PDF of

$$P[r] == \int_0^\infty \frac{\lambda^r e^{-\lambda}}{r!} f(\lambda)\, d\lambda == \frac{1}{r!} \int_0^\infty \lambda^r e^{-\lambda} f(\lambda)\, d\lambda \tag{1}$$

the PDF of a Log-normal distribution parametrized in terms of its actual mean and cv may be written as

$$\frac{e^{-\frac{\mathrm{Log}\left[\frac{\sqrt{1+c^2}\,\lambda}{\mu}\right]^2}{2\,\mathrm{Log}\left[1+c^2\right]}}}{\sqrt{2\pi}\,\lambda\,\sqrt{\mathrm{Log}\left[1+c^2\right]}}$$

Thus we may express the PDF of the PLN distribution as:

$$P[r] == \frac{1}{\sqrt{2\pi\,\mathrm{Log}\left[1+c^2\right]}\; r!} \int_0^\infty \lambda^{r-1} e^{-\lambda} e^{-\frac{\left(\mathrm{Log}\left[\lambda \frac{\sqrt{1+c^2}}{m}\right]\right)^2}{2\,\mathrm{Log}\left[1+c^2\right]}}\, d\lambda \tag{2}$$

Bullmer also shows, by calculate the factorial moments of this distribution by summing over r for fixed $\lambda$, and then integrating, that the k-th factorial moment of this distribution is equal to the k-th ordinary moment of the corresponding lognormal distribution, which in our parameterization is:

$$\left(1+c^2\right)^{\frac{k^2}{2}} \left(\frac{m}{\sqrt{1+c^2}}\right)^k$$

We note that the r-th raw moment of a random variable X can be expressed in terms of its factorial moments by the formula

$$\sum_{j=0}^{r} \left\{ \begin{matrix} r \\ j \end{matrix} \right\} E\left[(X)_j\right]$$

whereE$\left[(X)_j\right]$ is the $j$ − th factorial moment, and the curly braces denoteStirling numbers of the second kind, which are specified by the formula

$$\left\{ \begin{matrix} a \\ b \end{matrix} \right\} \to \frac{1}{b\,!} \sum_{i=0}^{b} (-1)^i \binom{b}{i} (b-i)^a$$

Putting this together gives us a formula for the raw moments of a Poisson log-normal distribution

$$r[k] = \sum_{j=0}^{k} \mathcal{S}_{k,j} \left(1 + c^2\right)^{\frac{j^2}{2}} \left( \frac{m}{\sqrt{1 + c^2}} \right)^j = \sum_{j=0}^{k} \mathcal{S}_{k,j} \left(1 + c^2\right)^{-\frac{j}{2} + \frac{j^2}{2}} m^j$$

where $\mathcal{S}_{k,j} = \left\{ \begin{matrix} k \\ j \end{matrix} \right\}$

Using this formula we may easily write the first five moments of the PLN distribution:

$r[1] = m,$

$r[2] = m + \left(1 + c^2\right) m^2,$

$r[3] = m + 3\left(1 + c^2\right) m^2 + \left(1 + c^2\right)^3 m^3,$

$r[4] = m + 7\left(1 + c^2\right) m^2 + 6\left(1 + c^2\right)^3 m^3 + \left(1 + c^2\right)^6 m^4,$

$r[5] = m + 15\left(1 + c^2\right) m^2 + 25\left(1 + c^2\right)^3 m^3 + 10\left(1 + c^2\right)^6 m^4 + \left(1 + c^2\right)^{10} m^5$

Note that when c<<1, this reduces to the known moments of the Poisson distribution, which are the Bell polynomials $B_k(m)$.

### Moments of modified, corrected Pearson residuals

The distribution of Pearson residuals under the null hypothesis will be the distribution of values of $\frac{(X-m)}{\alpha}$ where m and $\alpha$ are constants (in ordinary Pearson residuals, $\alpha = \sqrt{m}$; for modified corrected Pearson residuals $\alpha = \sqrt{m\left(1 + c^2\, m\right)}$). Division of the values of a distribution by any constant alters the moments of the distribution in a simple way: each moment *r[k]* is simply divided by $\alpha^k$.

The distribution of values *X-m*, where *X* is drawn from a PLN distribution, can be obtained from the definition of the *k*-th moment for the distribution of a random variable minus the constant *m*, which may be written as $\sum_{i=0}^{k} (x-m)^i\, P[x]\, dx$, where *P[x]* is the PDF of the random variable. Noting that $(x-m)^i$ can be expanded as $\sum_{i=0}^{k} \binom{k}{i} (-m)^{k-i}\, x^i$, we can re-write this as $\sum_{i=0}^{k} \binom{k}{i} (-m)^{k-i}\, x^i\, P[x]$ . Noting that $\sum_{i=0}^{k} x^i\, P[x]$ are simply the moments of the distribution defined by *P[x]*, we then get that the *k*-th moment of the distribution of *X-m* is simply

$$\sum_{i=0}^{k} \binom{k}{i} (-\mu \mathrm{m})^{k-i}\, r[i]$$

Accounting for division by $\alpha$, which in the case of modified corrected Pearson residuals will be $\sqrt{\mu\left(1 + c^2\, \mu\right)}$ gives us an equation for the k-th moments of a modified corrected Pearson residual, namely

$$\frac{1}{\left(\sqrt{m\left(1 + c^2\, m\right)}\right)^k} \sum_{i=0}^{k} \binom{k}{i} (-m)^{k-i}\, r[i]$$

where the *r[i]* are defined as above.

### Moments of modified corrected Fano factors

Modified corrected Fano factors ($\phi$') are obtained by summing over n squared modified corrected Pearson residuals, and dividing by n-1. Given the definition of the i-th moment of a distribution as the expectation values of $x^i$, it is easy to see that the moments of the square of a random variable are just the even-numbered moments of the random variable, i.e. moments 1,2,3... of squared Pearson residuals are simply the moments 2,4,6... of the Pearson residuals themselves. Accordingly, we replace the above formula with

$$\frac{1}{\left(m\left(1 + c^2\, m\right)\right)^k} \sum_{i=0}^{2k} \binom{2k}{i} (-m)^{2k-i}\, r[i]$$

Putting this together with the definitions of r[i] we may write the first 5 moments of the squared modified corrected Pearson residual, which we will denote as $\mathcal{E}_1$, as

$\mathcal{E}_1 \to 1,$

$$\mathcal{E}_2 \to \frac{1 + m\left(3 + c^2\left(7 + m\left(6 + 3\, c^2\, (6 + m) + c^4\left(6 + \left(16 + 15\, c^2 + 6\, c^4 + c^6\right) m\right)\right)\right)\right)}{m\left(1 + c^2\, m\right)^2},$$

$$\mathcal{E}_3 \to \frac{1}{m^2 (1 + c^2 m)^3} \Big(1 + 31 (1 + c^2) m + 90 (1 + c^2)^3 m^2 + 65 (1 + c^2)^6 m^3 + 15 (1 + c^2)^{10} m^4 - 5 m^5 + (1 + c^2)^{15} m^5 + 15 m^4 (1 + m + c^2 m) - 20 m^2 \big(m + 3 (1 + c^2) m^2 + (1 + c^2)^3 m^3\big) + 15 m \big(m + 7 (1 + c^2) m^2 + 6 (1 + c^2)^3 m^3 + (1 + c^2)^6 m^4\big) - 6 \big(m + 15 (1 + c^2) m^2 + 25 (1 + c^2)^3 m^3 + 10 (1 + c^2)^6 m^4 + (1 + c^2)^{10} m^5\big)\Big),$$

$$\mathcal{E}_4 \to \frac{1}{m^3 (1 + c^2 m)^4} \Big(1 + 127 (1 + c^2) m + 966 (1 + c^2)^3 m^2 + 1701 (1 + c^2)^6 m^3 + 1050 (1 + c^2)^{10} m^4 + 266 (1 + c^2)^{15} m^5 + 28 (1 + c^2)^{21} m^6 - 7 m^7 + (1 + c^2)^{28} m^7 + 28 m^6 (1 + m + c^2 m) - 56 m^4 \big(m + 3 (1 + c^2) m^2 + (1 + c^2)^3 m^3\big) + 70 m^3 \big(m + 7 (1 + c^2) m^2 + 6 (1 + c^2)^3 m^3 + (1 + c^2)^6 m^4\big) - 56 m^2 \big(m + 15 (1 + c^2) m^2 + 25 (1 + c^2)^3 m^3 + 10 (1 + c^2)^6 m^4 + (1 + c^2)^{10} m^5\big) + 28 m \big(m + 31 (1 + c^2) m^2 + 90 (1 + c^2)^3 m^3 + 65 (1 + c^2)^6 m^4 + 15 (1 + c^2)^{10} m^5 + (1 + c^2)^{15} m^6\big) - 8 \big(m + 63 (1 + c^2) m^2 + 301 (1 + c^2)^3 m^3 + 350 (1 + c^2)^6 m^4 + 140 (1 + c^2)^{10} m^5 + 21 (1 + c^2)^{15} m^6 + (1 + c^2)^{21} m^7\big)\Big),$$

$$\mathcal{E}_5 \to \frac{1}{m^4 (1 + c^2 m)^5} \Big(1 + 511 (1 + c^2) m + 9330 (1 + c^2)^3 m^2 + 34\,105 (1 + c^2)^6 m^3 + 42\,525 (1 + c^2)^{10} m^4 + 22\,827 (1 + c^2)^{15} m^5 + 5880 (1 + c^2)^{21} m^6 + 750 (1 + c^2)^{28} m^7 + 45 (1 + c^2)^{36} m^8 - 9 m^9 + (1 + c^2)^{45} m^9 + 45 m^8 (1 + m + c^2 m) - 120 m^6 \big(m + 3 (1 + c^2) m^2 + (1 + c^2)^3 m^3\big) + 210 m^5 \big(m + 7 (1 + c^2) m^2 + 6 (1 + c^2)^3 m^3 + (1 + c^2)^6 m^4\big) - 252 m^4 \big(m + 15 (1 + c^2) m^2 + 25 (1 + c^2)^3 m^3 + 10 (1 + c^2)^6 m^4 + (1 + c^2)^{10} m^5\big) + 210 m^3 \big(m + 31 (1 + c^2) m^2 + 90 (1 + c^2)^3 m^3 + 65 (1 + c^2)^6 m^4 + 15 (1 + c^2)^{10} m^5 + (1 + c^2)^{15} m^6\big) - 120 m^2 \big(m + 63 (1 + c^2) m^2 + 301 (1 + c^2)^3 m^3 + 350 (1 + c^2)^6 m^4 + 140 (1 + c^2)^{10} m^5 + 21 (1 + c^2)^{15} m^6 + (1 + c^2)^{21} m^7\big) + 45 m \big(m + 127 (1 + c^2) m^2 + 966 (1 + c^2)^3 m^3 + 1701 (1 + c^2)^6 m^4 + 1050 (1 + c^2)^{10} m^5 + 266 (1 + c^2)^{15} m^6 + 28 (1 + c^2)^{21} m^7 + (1 + c^2)^{28} m^8\big) - 10 \big(m + 255 (1 + c^2) m^2 + 3025 (1 + c^2)^3 m^3 + 7770 (1 + c^2)^6 m^4 + 6951 (1 + c^2)^{10} m^5 + 2646 (1 + c^2)^{15} m^6 + 462 (1 + c^2)^{21} m^7 + 36 (1 + c^2)^{28} m^8 + (1 + c^2)^{36} m^9\big)\Big),$$

From here we note that the Fano factor is obtained by taking a sum of squared, modified, corrected Pearson residuals, each of which is characterized by its own set of moments and its own value of m. Thus, we need add indices to the values of m, i.e. using m[j] to refer to the value of m in cell j.

To obtain the moments of the sum of random variables we use the fact that the moment generating function (MGF) of a sum of independent random variables is equal to the product of the MGFs of the individual random variables. In practice, it is easier to generate solutions by switching from moments to cumulants, for which an analogous rule exists, namely that the cumulant generating function (KGF) of a sum of independent random variables is the sum of the cumulant generating functions of the individual random variables. As the k-th cumulant is simply the k-th derivative of the KGF, and differentiation distributes over addition, we see that k-th cumulant of a sum of random variables is just the sum of the k-th cumulants of each of the random variables.

Cumulants may be obtained from moments by noting the the KGF is simply the logarithm of the MGF, from which the following relationship between the first 5 moments ($\mathcal{E}_1, \mathcal{E}_2, \mathcal{E}_3, \mathcal{E}_4, \mathcal{E}_5$) and first five cumulants ($\kappa_1, \kappa_2, \kappa_3, \kappa_4, \kappa_5$) may be derived

$$\kappa_1 = \mathcal{E}_1,$$
$$\kappa_2 = -\mathcal{E}_1^2 + \mathcal{E}_2,$$
$$\kappa_3 = 2 \mathcal{E}_1^3 - 3 \mathcal{E}_1 \mathcal{E}_2 + \mathcal{E}_3,$$
$$\kappa_4 = -6 \mathcal{E}_1^4 + 12 \mathcal{E}_1^2 \mathcal{E}_2 - 3 \mathcal{E}_2^2 - 4 \mathcal{E}_1 \mathcal{E}_3 + \mathcal{E}_4,$$
$$\kappa_5 = 24 \mathcal{E}_1^5 - 60 \mathcal{E}_1^3 \mathcal{E}_2 + 20 \mathcal{E}_1^2 \mathcal{E}_3 - 10 \mathcal{E}_2 \mathcal{E}_3 + 5 \mathcal{E}_1 \left(6 \mathcal{E}_2^2 - \mathcal{E}_4\right) + \mathcal{E}_5$$

Finally, recalling that the modified corrected Fano factor $\phi$' is obtained by summing over n squared, modified, corrected Pearson residuals and then dividing by n-1, to obtain the moments (or cumulants) of $\phi$' we must divide the k-th moment (or cumulant) by $(n-1)^k$. Doing this, a making the appropriate substitutions for $\kappa, \mathcal{E}$ and m in each cell, we obtain the following final formulae for the first five cumulants of $\phi$', where the terms have been made a bit more compact by substituting $\chi$ for $1 + c^2$

$$\kappa_1 \to \frac{n}{-1 + n},$$

$$\kappa_2 \to \frac{\sum_{j=1}^{n}\left(-1+\frac{1+m[j]\left(-4+7\chi+6\left(1-2\chi+\chi^3\right)m[j]+\left(-3+6\chi-4\chi^3+\chi^6\right)m[j]^2\right)}{m[j]\left(1+(-1+\chi)m[j]\right)^2}\right)}{(-1+n)^2},$$

$$\kappa_3 \to \frac{1}{(-1+n)^3}\left(\sum_{j=1}^{n}\frac{1}{m[j]^2\,(1+(-1+\chi)\,m[j])^3}\,\Big(1+(-9+31\chi)\,m[j]+2\left(16-57\chi+45\chi^3\right)m[j]^2+\right.$$
$$\left(-56+180\chi-21\chi^2-168\chi^3+65\chi^6\right)m[j]^3+3\left(16-48\chi+14\chi^2+40\chi^3-6\chi^4-21\chi^6+5\chi^{10}\right)m[j]^4+$$
$$\left.\left(-16+48\chi-24\chi^2-30\chi^3+12\chi^4+18\chi^6-3\chi^7-6\chi^{10}+\chi^{15}\right)m[j]^5\Big)\right),$$

$$\kappa_4 \to \frac{1}{(-1+n)^4}\left(\sum_{j=1}^{n}\frac{1}{m[j]^3\,(1+(-1+\chi)\,m[j])^4}\,\Big(1+(-15+127\chi)\,m[j]+\left(92-674\chi+966\chi^3\right)m[j]^2+\right.$$
$$\left(-302+1724\chi-271\chi^2-2804\chi^3+1701\chi^6\right)m[j]^3+6\left(96-452\chi+174\chi^2+620\chi^3-102\chi^4-511\chi^6+175\chi^{10}\right)m[j]^4+$$
$$2\left(-320+1344\chi-822\chi^2-1390\chi^3+672\chi^4+1124\chi^6-151\chi^7-590\chi^{10}+133\chi^{15}\right)m[j]^5+$$
$$4\left(96-384\chi+312\chi^2+278\chi^3-276\chi^4+18\chi^5-194\chi^6+84\chi^7-9\chi^9+126\chi^{10}-15\chi^{11}-43\chi^{15}+7\chi^{21}\right)m[j]^6+$$
$$\left(-96+384\chi-384\chi^2-160\chi^3+314\chi^4-48\chi^5+112\chi^6-120\chi^7+12\chi^8+24\chi^9-\right.$$
$$\left.\left.80\chi^{10}+24\chi^{11}-3\chi^{12}+32\chi^{15}-4\chi^{16}-8\chi^{21}+\chi^{28}\right)m[j]^7\Big)\right),\ \kappa_5 \to \frac{1}{(-1+n)^5}$$

$$\left(\sum_{j=1}^{n}\frac{1}{m[j]^4\,(1+(-1+\chi)\,m[j])^5}\,\Big(1+(-25+511\chi)\,m[j]+30\left(8-119\chi+311\chi^3\right)m[j]^2+5\left(-248+2540\chi-561\chi^2-7208\chi^3+6821\chi^6\right)\right.$$
$$m[j]^3+\left(3904-29\,880\chi+15\,690\chi^2+68\,000\chi^3-12\,990\chi^4-86\,865\chi^6+42\,525\chi^{10}\right)m[j]^4+$$
$$\left(-7872+49\,360\chi-39\,660\chi^2-81\,110\chi^3+46\,460\chi^4+98\,270\chi^6-13\,365\chi^7-74\,910\chi^{10}+22\,827\chi^{15}\right)m[j]^5+$$
$$20\left(512-2832\chi+2898\chi^2+3168\chi^3-3654\chi^4+216\chi^5-3109\chi^6+1499\chi^7-240\chi^9+2953\chi^{10}-315\chi^{11}-\right.$$
$$\left.1390\chi^{15}+294\chi^{21}\right)m[j]^6+10\left(-832+4288\chi-5136\chi^2-2940\chi^3+6222\chi^4-1008\chi^5+2276\chi^6-2778\chi^7+\right.$$
$$\left.172\chi^8+806\chi^9-2636\chi^{10}+842\chi^{11}-65\chi^{12}-90\chi^{13}+1420\chi^{15}-140\chi^{16}-476\chi^{21}+75\chi^{28}\right)m[j]^7+$$
$$5\left(768-3840\chi+5184\chi^2+1152\chi^3-5656\chi^4+1728\chi^5-936\chi^6+2432\chi^7-420\chi^8-960\chi^9+1448\chi^{10}-\right.$$
$$\left.912\chi^{11}+186\chi^{12}+192\chi^{13}-720\chi^{15}+170\chi^{16}-12\chi^{18}+288\chi^{21}-28\chi^{22}-73\chi^{28}+9\chi^{36}\right)m[j]^8+$$
$$\left(-768+3840\chi-5760\chi^2+320\chi^3+5280\chi^4-2536\chi^5+560\chi^6-2240\chi^7+840\chi^8+980\chi^9-1072\chi^{10}+880\chi^{11}-\right.$$
$$\left.\left.300\chi^{12}-210\chi^{13}+400\chi^{15}-180\chi^{16}+20\chi^{17}+40\chi^{18}-170\chi^{21}+40\chi^{22}+50\chi^{28}-5\chi^{29}-10\chi^{36}+\chi^{45}\right)m[j]^9\Big)\right)$$

### Moments of modified corrected PCCs

The procedure for deriving the moments, and subsequently cumulants, of PCC’ are similar to those used above for $\phi$’, except that certain approximations are required due to the more complicated set of terms that go into defining PCC’. As described in the main text, the PCC is fundamentally an average over products of pairs of Pearson residuals, i.e.

$$\text{PCC'} = \frac{1}{(n-1)\,\sqrt{\phi'_a\,\phi'_b}}\sum_{j=1}^{n} P'_{a,j}\,P'_{b,j}$$

where $P'_{a,j}$ and $P'_{b,j}$ represent the modified corrected Pearson residuals for genes a and b, respectively, in cell j. Calculating the moments of $\sum_{j=1}^{n} P'_{a,j}\,P'_{b,j}$ requires only one additional rule beyond those enumerated in the previous section, which is that that moment of the product of two random variables is the product of the moments of the individual random variables. With this information it is possible to write out a general formula for the moments of $\sum_{j=1}^{n} P'_{a,j}\,P'_{b,j}$ .

However, accounting for the further division by $(n-1)\,\sqrt{\phi'_a\,\phi'_b}$ is less straightforward because $\phi'_a$ and $\phi'_b$ are not constants, but are rather distributed according to the Fano factor distributions of genes a and b, which we had just calculated.

Moreover, not only must $\phi'_a$ and $\phi'_b$ be treated as random variables, they are technically not independent of the other random variables, i.e. those that define the modified corrected Pearson residuals. Despite this, however, when the number of cells is in the hundreds or greater, they may be treated as *very nearly* independent of the individual modified corrected Pearson residuals because they effectively average over a very large number of those residuals.

Doing that leaves us only to calculate the moments of $1 / \sqrt{\phi'}$ for each gene, which we may then multiply by the moments of $\frac{1}{(n-1)} \sum_{j=1}^{n} P'_{a,j} P'_{b,j}$ to produce the moments of PCC'. Since the first moment of $\frac{1}{(n-1)} \sum_{j=1}^{n} P'_{a,j} P'_{b,j}$ under the null hypothesis is zero (no correlation), we don't actually need the first moments of $1 / \sqrt{\phi'}$, just the higher ones.

Unfortunately, analytically deriving the moments for the square root of a random variable with an arbitrary distribution is not generally straightforward, nor is the derivation of moments of the multiplicative inverse of such a distribution. As it happens, when the number of cells is large (at least many hundreds), these moments turn out to be rather close to 1, i.e. simply omitting them has relatively little effect on the moments of PCC'. However, since we cannot always be certain that this will be the case, we developed an approach to approximate their values by simulation.

Briefly, for any given dataset with *m* genes and *n* cells, for which a value of the coefficient of variation of gene expression noise has been chosen to be *c*, we first use the cell specific sequencing depths to calculate the rows of the expectation matrix that would be generated for 8 values of total expression (sum over the cells of the UMI for any given gene) ranging from the lowest to the highest in the data set (and distributing them logarithmically). From each row of this 8 x *n* matrix, we repeatedly generate a vector by replacing each entry $\mu_i$ with a Poisson Random Variate or a Random Variate of a Log Normal distribution with mean $\mu_i$ and coefficient of variation *c*. We then calculate a modified corrected Fano factor using these entries and takes its square root and multiplicative inverse. We gather a sufficiently large number of these that the first 5 moments of their distribution adequately converge; not surprisingly this requires a larger number of such vectors when the total gene count is low. Typically, between 300 and 2000 iterations are required, depending on the level of gene expression.

In this way we generate empirical estimates of the moments of $1 / \sqrt{\phi'}$ for 8 representative levels of gene expression. From there it is straightforward to use interpolation to estimate the moments of $1 / \sqrt{\phi'}$ for any level of gene expression in our data set. Finally to obtain the moments of $1/\sqrt{\phi'_a \phi'_b}$ for pairs of genes a and b we simply multiply the moments of $1/\sqrt{\phi'_a}$ and $1/\sqrt{\phi'_b}$ (since they are independent).

All told, given that the k-th moment of any constant $\varphi$ times a random variable is $\varphi^k$ times the k-th moment of the random variable, we get that, for PCC':

$$\mathcal{E}_k(\mathrm{PCC}_{a,b}) = \mathcal{E}_k\left(\frac{1}{(n-1)\sqrt{\phi'_a \phi'_b}} \sum_{j=1}^{n} P'_{a,j} P'_{b,j}\right) = \frac{1}{(n-1)^k} \mathcal{E}_k\left(\frac{1}{\sqrt{\phi_a}}\right) \mathcal{E}_k\left(\frac{1}{\sqrt{\phi_b}}\right) \mathcal{E}_k\left(\sum_{j=1}^{n} P'_{a,j} P'_{b,j}\right)$$

where $\mathcal{E}_k$ stands for the k-the moment, and a and b are individual genes. Converting the raw moments to cumulants gives the following formula, where $\kappa_k$ representes the $k-$th cumulant, and $m_a[j]$ and $m_b[j]$ stand for the values in column j of the expectation matrix for genes a and b, respectively.

$$\kappa_1 \to 0;$$

$$\kappa_2 \to \frac{n}{(n-1)^2} \mathcal{E}_2\left(\frac{1}{\sqrt{\phi_a}}\right) \mathcal{E}_2\left(\frac{1}{\sqrt{\phi_b}}\right);$$

$$\kappa_3 \to \frac{1}{(n-1)^3} \mathcal{E}_3\left(\frac{1}{\sqrt{\phi_a}}\right) \mathcal{E}_3\left(\frac{1}{\sqrt{\phi_b}}\right) \sum_{j=1}^{n} \frac{m_a[j]\left(1+3\,c^2\,m_a[j]+c^4\left(3+c^2\right)m_a[j]^2\right) m_b[j]\left(1+3\,c^2\,m_b[j]+c^4\left(3+c^2\right)m_b[j]^2\right)}{\left(m_a[j]\left(1+c^2\,m_a[j]\right)\right)^{3/2}\left(m_b[j]\left(1+c^2\,m_b[j]\right)\right)^{3/2}};$$

$$\kappa_4 \to \frac{1}{(n-1)^4} \left(-3\,n\left(\mathcal{E}_2\left(\frac{1}{\sqrt{\phi_a}}\right)\mathcal{E}_2\left(\frac{1}{\sqrt{\phi_b}}\right)\right)^2 + \mathcal{E}_4\left(\frac{1}{\sqrt{\phi_a}}\right)\mathcal{E}_4\left(\frac{1}{\sqrt{\phi_b}}\right) \sum_{j=1}^{n} \left(\left(1 + \left(3 + 7\,c^2\right)m_a[j] + 6\left(c^2 + 3\,c^4 + c^6\right)m_a[j]^2 + c^4\left(3 + 16\,c^2 + 15\,c^4 + 6\,c^6 + c^8\right)m_a[j]^3\right)\left(1 + \left(3 + 7\,c^2\right)m_b[j] + 6\left(c^2 + 3\,c^4 + c^6\right)m_b[j]^2 + c^4\left(3 + 16\,c^2 + 15\,c^4 + 6\,c^6 + c^8\right)m_b[j]^3\right)\right) \Big/ \left(m_a[j]\left(1 + c^2\,m_a[j]\right)^2 m_b[j]\left(1 + c^2\,m_b[j]\right)^2\right)\right);$$

$$\kappa_5 \to \frac{1}{(n-1)^5} \left(-10\,\mathcal{E}_2\left(\frac{1}{\sqrt{\phi_a}}\right)\mathcal{E}_2\left(\frac{1}{\sqrt{\phi_b}}\right)\mathcal{E}_3\left(\frac{1}{\sqrt{\phi_a}}\right)\mathcal{E}_3\left(\frac{1}{\sqrt{\phi_b}}\right) \sum_{j=1}^{n} \left[\frac{1+c^2\,m_a[j]\left(3+c^2\left(3+c^2\right)m_a[j]\right)}{\sqrt{m_a[j]}\left(1+c^2\,m_a[j]\right)^{3/2}} * \frac{1+c^2\,m_b[j]\left(3+c^2\left(3+c^2\right)m_b[j]\right)}{\sqrt{m_b[j]}\left(1+c^2\,m_b[j]\right)^{3/2}}\right] + \mathcal{E}_5\left(\frac{1}{\sqrt{\phi_a}}\right)\mathcal{E}_5\left(\frac{1}{\sqrt{\phi_b}}\right) \sum_{j=1}^{n} \frac{1}{m_a[j]^{3/2}\left(1+c^2\,m_a[j]\right)^{5/2} m_b[j]^{3/2}\left(1+c^2\,m_b[j]\right)^{5/2}} \left(1 + 5\left(2 + 3\,c^2\right)m_a[j] + 5\,c^2\left(8 + 15\,c^2 + 5\,c^4\right)m_a[j]^2 + 10\,c^4\left(6 + 17\,c^2 + 15\,c^4 + 6\,c^6 + c^8\right)m_a[j]^3 + c^6\left(30 + 135\,c^2 + 222\,c^4 + 205\,c^6 + 120\,c^8 + 45\,c^{10} + 10\,c^{12} + c^{14}\right)m_a[j]^4\right) \left(1 + 5\left(2 + 3\,c^2\right)m_b[j] + 5\,c^2\left(8 + 15\,c^2 + 5\,c^4\right)m_b[j]^2 + 10\,c^4\left(6 + 17\,c^2 + 15\,c^4 + 6\,c^6 + c^8\right)m_b[j]^3 + c^6\left(30 + 135\,c^2 + 222\,c^4 + 205\,c^6 + 120\,c^8 + 45\,c^{10} + 10\,c^{12} + c^{14}\right)\right);$$

### From cumulants and observations to *p*-values

Although there is no analytical method for reconstructing an arbitrary distribution from a finite number of its moments, several methods exist for approximating the body or tails of such distributions. Since we are interested in assessing the probabilities of relatively rare events, we need a way to estimate tail probabilities reasonably accurately. The Cornish Fisher expansion [Cornish and Fisher, 1938] provides a useful algorithm for doing so. Briefly, from a finite number of cumulants of the distribution under the null hypothesis, and an observed value (in this case an observed PCC'), this expansion estimates the probability of that observation occurring under the null hypothesis, i.e. a *p*-value. Doing so requires finding the closest real root to zero of polynomials whose coefficients are derived from the cumulants and the observed data. We solve one such equation for each PCC', using a variety of shortcuts to speed the process. Given the existence of potential error in the estimation of moments and PCC' values from finite numbers of cells (see Sources of Error, below), it is sometimes necessary to accept roots when they bring the polynomial close to, but not exactly equal to, zero. This is particularly important when both of the genes involved are of very low expression level, such that the data vectors are very sparse.

### Estimating False Discovery rate

The Bonferroni correction is much too conservative for use in multiple hypothesis testing correction because pairwise correlations are far from independent measurements. We find that the Benjamini Hochberg procedure [Benjamini and Hochberg, 1995] for false-discovery rate estimation works reasonably well.

### Sources of Error

The values in the expectation matrix are estimates based on a finite sample, not the true mean for gene expression. The error introduced by this is discussed by Lause et al., (2021). It is unlikely to create a significant problem for genes where the total UMI is relatively large (i.e. where the number of cells times the observed UMI per cell is above a few hundred). However, since it is the total number of observations, and not the observations per cell, that dictate the uncertainty in sample estimates, we may expect that the fewer the cells, and the more shallow their sequencing, the greater the magnitude of this problem will be. In practice we find that, with typical sequencing depths, working with samples of at least 500 cells is advisable.

For genes in which total UMI is small, as mentioned above, the errors in estimation can be sufficient to shift the Cornish Fisher polynomial equations so that curves that should have crossed the abscissa near the origin (indicative of a high *p*-value) may fail to do so, curving away and returning to cross it elsewhere; this typically produces a highly erroneous *p*-value, which can usually be spotted on inspection of the raw gene expression vectors. To eliminate such cases, in all instances where the two genes being tested for correlation are both expressed at very low levels (a threshold of 84 total UMI across all cells is used), Cornish Fisher polynomials are tested for the presence of an extremum near the origin, and those gene-pairs are rejected if they display one.

In BigSur the distribution we use for gene expression noise is log-normal; furthermore the value of c for that distribution is taken to be constant for all genes. While theoretical arguments and empirical data support the use of the lognormal distribution [Bahar Halpern, 2015; Beal 2017], some theoretical arguments suggest that different distributions may be more appropriate for some genes, but it seems unlikely that the precise choice of distribution matters greatly to the results obtained by BigSur. Of greater consequence is the assumption of a common value of c, the coefficient of variation of this distribution, for every gene, when the value of c is likely to be different for different genes. In future, with greater knowledge, it may be possible to introduce gene-specific corrections to c. In the meantime, it is helpful to know that the choice of c only impacts the statistics for genes with relatively high expression level, among which correlations are most easily detected.

In determining the moments of PCC' under the null hypothesis, we use moment values for the inverse of the square root of the modified corrected Fano factors that are calculated by simulation and interpolation, using the actual sequencing depth distribution among the cells in the data set. As the simulations involve finite samples, there will be a small amount of error in these values. Furthermore, when we obtain the moments of PCC' by multiplying these moments by the moments of the sums of products of pairs of Pearson residuals, we are implicitly assuming that the Fano factors are independent of the distribution of the products of pairs of Pearson residuals. While technically this is not the case, the fact that the Fano factors average over a very large number of (squared) Pearson residuals makes them effectively independent. When cell numbers are small, however, this may cease to be true (providing a second reason not to apply BigSur to datasets with small numbers of cells). As discussed above, the moments of the inverse of the square root of the modified corrected Fano factors generally turn out to be quite close to 1, unless gene expression levels are low; in practice this means that correlations found to be significant when these moments are ignored tend to be almost identical to those obtained when they are included, suggesting that accuracy in determining their values is not very important.
